# Defining totipotency using criteria of increasing stringency

**DOI:** 10.1101/2020.03.02.972893

**Authors:** Eszter Posfai, John Paul Schell, Adrian Janiszewski, Isidora Rovic, Alexander Murray, Brian Bradshaw, Tine Pardon, Mouna El Bakkali, Irene Talon, Natalie De Geest, Pankaj Kumar, San Kit To, Sophie Petropoulos, Andrea Jurisicova, Vincent Pasque, Fredrik Lanner, Janet Rossant

**Author notes:** authors contributed equally (Eszter Posfai, John Paul Schell, Adrian Janiszewski).

## Abstract

Totipotency is the ability of a single cell to give rise to all the differentiated cells that build the conceptus, yet how to capture this property *in vitro* remains incompletely understood. Defining totipotency relies upon a variety of assays of variable stringency. Here we describe criteria to define totipotency. We illustrate how distinct criteria of increasing stringency can be used to judge totipotency by evaluating candidate totipotent cell types in the mouse, including early blastomeres and expanded or extended pluripotent stem cells. Our data challenge the notion that expanded or extended pluripotent states harbor increased totipotent potential relative to conventional embryonic stem cells under *in vivo* conditions.

## Introduction

During early mammalian development, the totipotent state of early blastomeres is rapidly lost as cells gradually restrict their developmental potential and commit to distinct cell lineages by the blastocyst stage^1,2,3^. In mouse, the first cell fate decision starting at embryonic day (E)2.5 sets aside the trophectoderm (TE), the precursors of the placenta, from the inner cell mass (ICM). A second cell fate decision starting around E3.5 within the ICM gives rise to the pluripotent epiblast (EPI) and the primitive endoderm (PE), precursors of all embryonic germ layers and extraembryonic yolk sac, respectively^4^. For the most part EPI, PE and TE cells maintain blastocyst-defined lineage assignments throughout subsequent development, with a notable exception for the PE, which was shown to also contribute to otherwise EPI-derived definitive endodermal lineages during post-implantation stages^5,6^.

While EPI, PE and TE cells exist only transiently in the developing embryo, distinct self-renewing stem cell types can be derived from each cell type using a combination of appropriate growth factors and/or inhibitors which capture and preserve their developmental potential in culture^7,8,9,10^. Murine embryonic stem cells (ESCs) derived from the early EPI lineage were originally established using fetal bovine serum and leukemia inhibitory factor (Lif)^7,8^. They can however also be cultured under naïve conditions, using inhibitors against mitogen-activated protein kinase and glycogen synthase kinase-3, termed 2i, in combination with Lif^11^. Trophoblast stem cells (TSCs) can be derived from the TE lineage using fibroblast growth factor 4 and heparin^9^ and extraembryonic endoderm (XEN) stem cells can be established from the PE using various methods^10,12,13^. Importantly, while each of these stem cell types are able to re-enter the normal course of embryonic development and differentiate into the downstream cell types similar to their *in vivo* counterparts, they are also lineage restricted in that they do not readily cross lineage boundaries that have been set during blastocyst formation^14,15^.

Strict lineage restriction differs between the three stem cell types, and is reflected in the time elapsed since the source lineages parted ways during embryo development. The closer relationship between EPI and PE lineages is also underscored by the observation that XEN cells can spontaneously appear in ESC cultures^16,17^ and ESCs can be readily converted into XEN cells using only soluble factors^13^. On the other hand, ESCs have only been reported to rarely contribute to trophectoderm-derived lineages *in vivo*^18^. Work by several laboratories has also shown that stem cell types with properties of TE, EPI or PE can be obtained by reprogramming lineage restriction using transcription factor (TF) expression. Long-term TF overexpression was shown to reprogram ESCs into TSC-like cells *in vitro*^19,20,21,22^. Induced pluripotent stem cells (iPSCs), as well as induced TSCs and induced XEN stem cells have also been derived by TF overexpression followed by culture with the appropriate growth media^23,24^. Additionally, mouse primed epiblast stem cells (EpiSCs), isolated from the post-implantation epiblast^25,26^ can be reverted back into ESCs^27^. Collectively, these studies suggested that it might be possible to induce totipotent stem cells, or at least cells that approach the totipotent state, by conversion from pluripotent stem cells.

In the past years there has been several reports of conditions to derive novel mouse stem cells types with the ability to produce descendants contributing to all three blastocyst-defined lineages^28,29,30,31,32,33,34^. In particular, two methods were described to derive extended or expanded pluripotent stem cells (EPSCs) by conversion from pre-existing ESCs or directly from 8-cell stage blastomeres^33,32^. In the first method, Liu lab EPSCs (L-EPSCs)^32^ were derived in Lif, CHIR, PD0325901, JNK Inhibitor VIII, SB203580, A-419259 and XAV939. In the second method, Deng lab EPSCs (D-EPSCs)^33^ were derived using Lif, CHIR, DiM ((S)-(+)-Dimethindene maleate), and MiH (Minocycline hydrochloride). Both cell types showed molecular and functional features that suggested expanded pluripotency, such as totipotency-associated marker gene expression and contribution to the EPI, PE as well as TE lineages using chimeric assays. Additionally, recent studies reported the ability of EPSCs, alone or in combination with TSCs, to self-assemble into blastocyst-like structures, termed blastoids, that contain cells with features of all three embryonic lineages^35,36^. These studies suggested that stem cells with the potential to give rise to both ICM and TE lineages, properties that define totipotent stem cells, can be isolated and expanded *in vitro*.

Many criteria of variable stringency can be used to assess totipotency. One criterion is to assess gene expression, in search of activated totipotency-associated marker genes. This can either be performed in bulk for a set of genes or through a more stringent approach taking advantage of transcriptome-wide single cell correlation analysis with totipotent cells of early embryos. More demanding is providing evidence of the potential to enter both the embryonic and extraembryonic pathway using *in vitro* differentiation assays. Finally, a more stringent requirement for evaluating the potential of different stem cell types is to perform *in vivo* aggregation experiments, by combining candidate cells with a host embryo and analyzing lineage contributions in the resulting chimera. Candidate cells are typically combined with morula (8-16 cell stage) or blastocyst stage host embryos and analyzed at different developmental stages. It is important to analyze chimeric contributions not only based on localization, but also by assessing lineage integration using functional marker analysis.

Here we subject candidate totipotent stem cells to these assays of increasing stringency to assess their developmental potential. We analyze the transcriptome and gene regulatory networks of ESCs, L-EPSC and D-EPSCs and pre-implantation embryos using bulk and single-cell RNA-sequencing (RNA-seq), and provide a resource for the community enabling interactive online data exploration. We investigate the ability of EPSCs to give rise to TSCs in both a conversion and a reprogramming setting. We analyze the transcriptome and gene regulatory networks of blastoids derived from EPSCs. Finally, we examine how EPSCs and blastomeres perform in chimeric experiments. We present a gold standard for analyzing contribution to different lineages, with a focus on contribution to the trophoblast lineage at different stages combined with molecular analyses. We emphasize the importance of thorough analysis of cell potential using high stringency assays and highlight the ongoing challenges of unlocking the totipotent state.

## Results

### Transcriptional signatures of preimplantation embryos and different stem cell states

Transcriptomic analysis can serve as effective means to monitor cellular states and analyze marker gene expression. Therefore, using transcriptional similarity analysis, we investigated which *in vivo* developmental stage or previously established *in vitro* stem cell state L-EPSCs and D-EPSCs resemble the most. First, we converted naïve ESCs (2iLif) to L-EPSCs and D-EPSCs using published protocols^33,32^. We observed similar morphological changes after conversion as previously reported^33,32^ and were able to stably maintain L-EPSC and D-EPSC cell lines (Figure S1A). In our first experiments we used bulk RNA sequencing for genome-wide detection of transcription and assessment of totipotency marker gene expression in L-EPSCs. We first set out to explore the dynamics with which a transcriptome shift is induced after switching ESCs into L-EPSC conditions (Figure 1A). Our results reveal a rapid transcriptome change, within 3 days of induction, indicating that ESCs can readily convert into L-EPSCs (Figure 1B). Intriguingly, despite these differences between L-EPSCs and ESCs, the L-EPSC transcriptome resembled the ESC transcriptome more than any early mouse embryo stage (Figure 1C and S2A and B) and 4-cell and 8-16 cell stage embryo marker genes remained mostly silenced (Figure 1D).

**Fig. 1:**
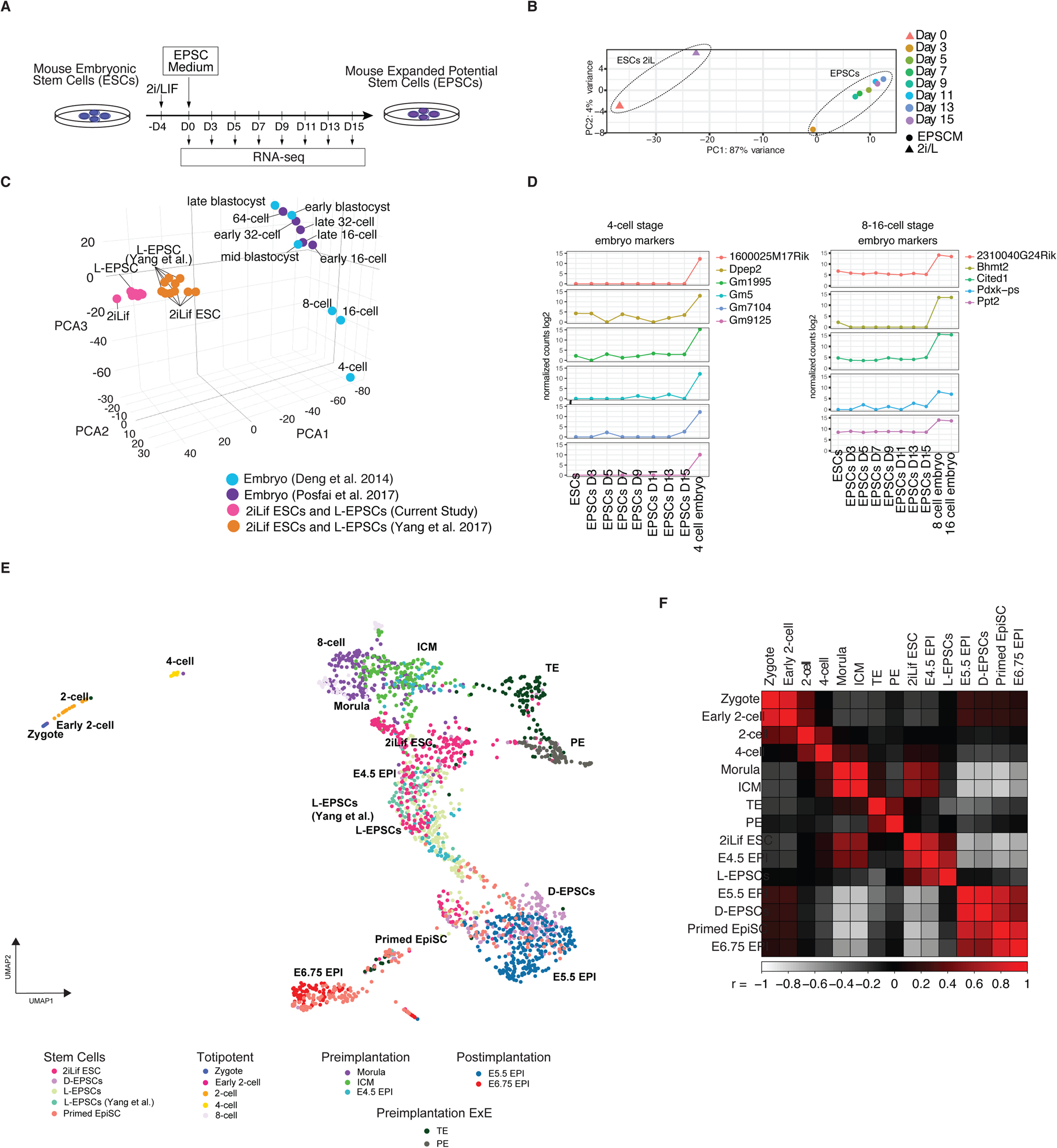
Gene expression analysis of candidate totipotent stem cells using RNA-seq. A) Experimental design of L-EPSC conversion. Mouse ESCs grown in 2iLif were switched to L-EPSC medium and subjected to bulk RNA-seq at day (D) 0, D3, D5, D7, D9, D11, D13 and D15. B) The L-EPSC transcriptome was rapidly induced during conversion from 2iLif ESCs. Principal Component Analysis (PCA) analysis of bulk RNA-seq data of samples from different timepoints of L-EPSC conversion and 2iLif ESCs. C) PCA analysis of bulk L-EPSC transcriptomes after integrating the dataset from this study with published scRNA-seq data (Deng *et al*. 2014^37^; Posfai *et al*. 2017^3^; Yang *et al*. 2017^32^). D) Silencing of most 4-cell, 8-cell and 16-cell stage marker genes is maintained in L-EPSCs. Expression of 4-cell and 8-16 cell stage marker genes in ESCs undergoing conversion to L-EPSCs. E) Single-cell UMAP comparing developmental progression from zygote to E6.75 EPI, including preimplantation extra embryonic lineages, with L-EPSCs, previously published L-EPSCs^32^, D-EPSCs, 2iLif ESCs and primed EpiSCs. F) Correlation matrix based on top 2000 expressed genes averaged tracking zygote to E6.75 EPI, including preimplantation extraembryonic lineages, with L-EPSCs, D-EPSCs, 2iLif ESCs and primed EpiSCs.

Single-cell RNA-seq (scRNA-seq) is particularly suited to resolve cellular heterogeneity and identify subpopulations with distinct transcriptional features. To examine whether totipotent features can be detected in individual cells, we applied SMART-seq2 scRNA-seq to ESCs, as well as to L-EPSCs and D-EPSCs derived from them. As a reference we transcriptionally tracked mouse preimplantation lineage segregation and post-implantation epiblast development from zygote to E6.75, and included naïve ESC and primed EpiSC states as well (Figure 1E). This dataset includes single-cell sequencing data from Deng *et al*. 2014^37^, Posfai *et al*. 2017^3^, Mohammed *et al*. 2017^38^, and Chen *et al*. 2016^39^, as well as an additional 96 cells from E2.5 and E4.5 embryos, and 551 cells from three pluripotent stem cell conditions: ESCs cultured in 2iLif, L-EPSCs and D-EPSCs, sequenced in this study. This integrated dataset provided suitable sampling for establishing a comparison of embryonic development with the different stem cell culture conditions, using Seurat v3.0 CCA integration tools (Figure S2C). Based on clustering using the top 2000 most variable genes, we resolved clear segregation of ICM/TE and EPI/PE lineages and annotated them based on the expression of well-established lineage markers (Figure S2D). As expected, ESCs cultured in 2iLif conditions occupied the space between E3.5 ICM and E4.5 EPI, while primed EpiSCs clustered with E5.5 and E6.75 EPI cells. We found that along the embryonic developmental trajectory L-EPSCs, as parental ESCs, clustered between the E3.5 ICM and E4.5 EPI stages, while the majority of D-EPSCs clustered together with E5.5 stage EPI cells (Figure 1E). We observed that top differentially expressed genes reported to be upregulated in D-EPSCs compared to Lif/serum-cultured ESCs were also upregulated in D-EPSCs compared to 2iLif-cultured ESCs (Figure S2E). We additionally constructed a correlation matrix from the top 2000 genes averaged expression to compare each stem cell condition independently with each developmental stage (Figure 1F). While ESCs showed high correlation with all preimplantation stages: 8-cell (r = .45, p < .001), morula (r = .55, p < .001), ICM (r = .53, p < .001), and highest similarity with E4.5 EPI (r = .72, p < .001), L-EPSCs (from both this and previous study) exhibited the most resemblance to E4.5 EPI (r = .64, p < .001), while lacking significant correlation with other developmental stages (p > .05). Consistent with the UMAP, D-EPSCs correlated the most with E5.5 EPI (r = .89, p < .001), but also showed close similarity with primed EpiSCs (r = .71, p < .001), and E6.75 EPI (r = .51, p < .001). The position occupied by L-EPSCs in the UMAP space is consistent with the original report by Yang *et al*.^32^. In conclusion, L- and D-EPSCs single-cell transcriptomes align with pluripotent rather than totipotent states.

Embryo development is under the control of transcription factors that bind to cis-regulatory regions, forming gene regulatory networks. We reconstructed gene regulatory networks which are active in early development, ESCs and EPSCs, from scRNA-seq data, using single-cell gene regulatory inference and clustering (pySCENIC^40^). SCENIC predicts TFs that may control cellular states present in the dataset, together with candidate TF target genes. A TF and its candidate targets are called a regulon, and by quantifying the activity of regulons in each single cell, SCENIC can be used to cluster cells based on the activity of regulatory programs. In contrast to identifying the global transcriptional state of cells, here we project a UMAP visualization based on regulon activity and reveal that 2iLif ESCs localize closest to E3.5 ICM, and both L- and D-EPSCs clustered between E4.5 EPI and E5.5 EPI (Figure S2F). These results show that the regulatory state of EPSCs resembles that of late pluripotent EPI rather than earlier, totipotent developmental stages.

### Capacity of EPSCs to enter the trophectoderm program and generate TSC-like cells *in vitro*

Another test to judge totipotency is to evaluate the capacity of cells to enter the trophoblast linage. This can be assessed *in vitro* by switching cells to TSC culture conditions and assaying whether cells give rise to TSC-like cells, a transition that pluripotent ESCs cannot make. When bulk L-EPSCs cultures were switched to TSC conditions followed by RT-PCR to assay trophoblast marker gene expression (Figure 2A), no substantial activation of such genes was seen (Figure 2B).

**Fig. 2:**
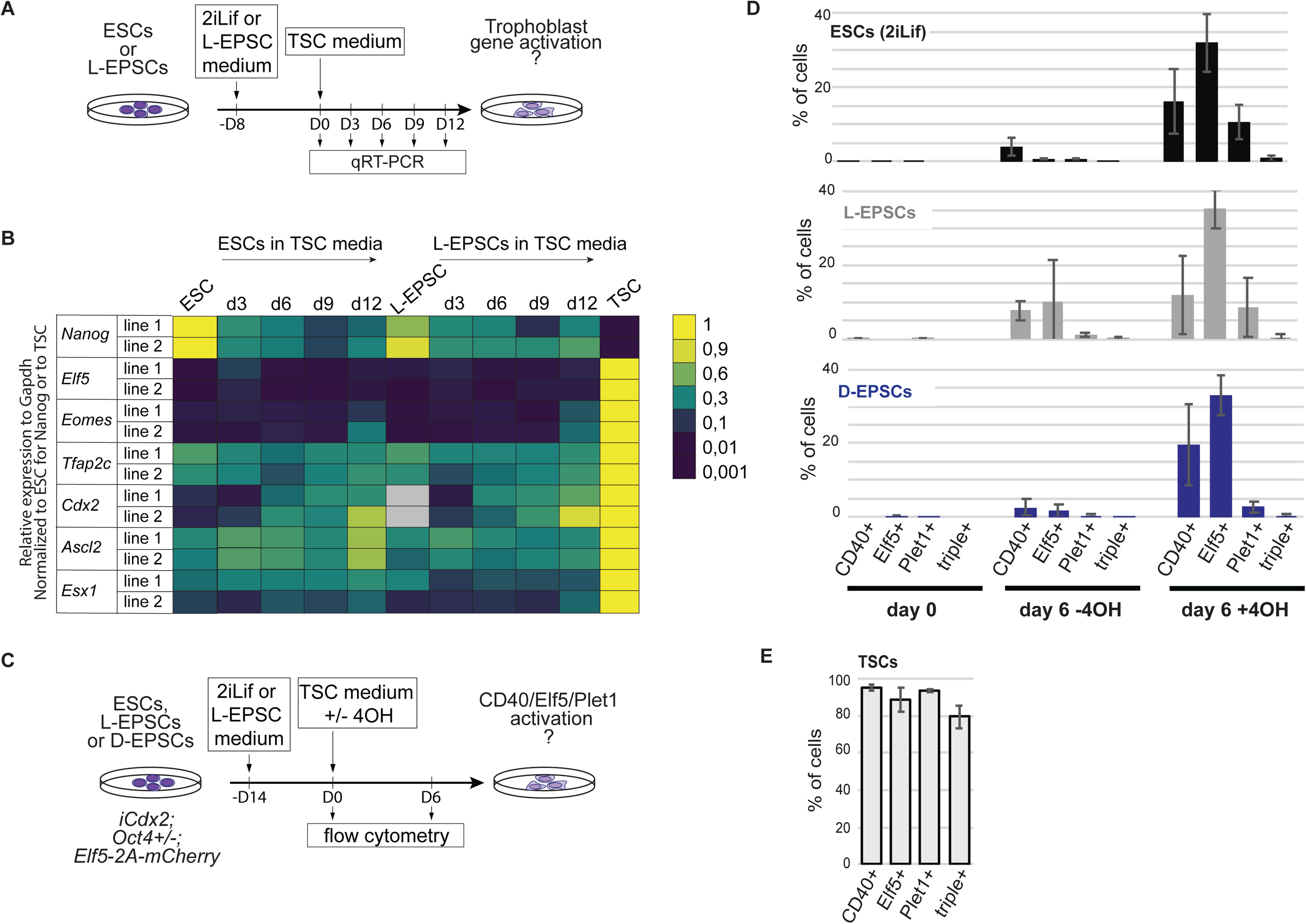
*In vitro* capacity and potential of candidate totipotent stem cells to give rise to trophoblast stem cells. A) Differentiation of L-EPSCs to TSCs. 2iLif ESCs or L-EPSCs were subjected to TSC differentiation conditions. RNA was collected at D0, D3, D6, D9, D12 and qPCR was used to measure the expression of TE marker genes. B) Silencing of extra-embryonic gene expression was maintained in L-EPSCs exposed to TSC differentiation conditions. Expression of TSC markers: *Elf5, Eomes, Tfap2c, Cdx2, Ascl2, Esx1* and *Nanog* as control at D3, D6, D9 and D12 of differentiation by RT-qPCR. C) Experimental design of ESC, L-EPSC or D-EPSC to TSC conversion (−4OH) or reprogramming experiments. ESC, L-EPSC or D-EPSC lines harbored a tamoxifen(4OH)-inducible Cdx2 transgene, were heterozygous for Oct4 (*Pou5f1*) and contained an Elf5-2a-mCherry reporter. Cells switched to TSC media with or without 4OH were analyzed by flow cytometry on indicated days for TSC markers CD40, ELF5 and PLET1. D) Flow cytometry analysis of ESC, L-EPSC or D-EPSC cells switched to TSC media with or without 4OH on indicated days. Percentages of cells positive of each TSC marker, or cells positive for all three markers are shown. Three biological replicates were performed, error bars indicate standard deviation of mean. E) Flow cytometry analysis of TSCs containing an Elf5-2s-mCherry reporter. Percentages of cells positive of each TSC marker, or cells positive for all three markers are shown. Three biological replicates were performed, error bars indicate standard deviation of mean.

To examine whether a small subpopulation of L-EPSCs or D-EPSCs may harbor the potential to directly convert into TSCs, as suggested by a previous study (Yang^32^), we analyzed the expression of key TSC markers on a single cell level using flow cytometry (Figure 2C). Additionally, we tested whether L-EPSCs and D-EPSCs can be reprogrammed into TSC-like cells more efficiently than ESCs. While ESCs do not readily convert into TSCs, they can be reprogrammed with low efficiency into TSCs by induced overexpression of TSC-associated TFs, such as *Cdx2* and by lowering the expression of the pluripotency factor Oct4^19^. To assay TSC reprogramming, we used a tetracycline-inducible *Cdx2* (iCdx2) and Oct4 heterozygous ESC background, used in the original ESC-to-TSC reprogramming experiments^19^. To read out fate conversion using flow cytometry, we immunostained for two TSC-specific cell surface markers, CD40 and Plet1. We also established an Elf5-2A-mCherry reporter ESC line by targeting 2A-mCherry to the C-terminus of the endogenous *Elf5* locus^41,42,20,22^ (Figure 2C). We then switched L-EPSCs, D-EPSCs and ESCs to TSC medium (with Fgf4 and Heparin) with or without tetraycycline (+/-4OH) and cultured the cells for 6 days (6d) before analyzing TSC-marker expression. We found that both in the absence or presence of Cdx2 induction, there were no significant differences in the number of single or triple marker positive cells between ESC and either EPSC conditions (Figure 2D). In contrast, a control TSC line in which we also targeted the Elf5 gene with the mCherry reporter showed 80% CD40/PLET1/ELF5 triple-positive cells (Figure 2E). Collectively, these results indicate that the EPSC states do not facilitate more efficient reprogramming into TSCs *in vitro*, in contrast with a previous study that suggested an increased ability of EPSC to give rise to TSC-like cells compared to ESC^33,32^.

### *In vitro* blastoid-forming ability of D-EPSCs based on Li *et al*. 2019 and Sozen *et al*. 2019

The ultimate proof of totipotency is the ability of a single cell type, or more stringently a single cell, to give rise to an entire blastocyst and subsequently a viable and fertile animal. Recently, blastocyst-like structures, termed blastoids, have been generated *in vitro* from different stem cell types^43,44,35,36^. These protocols use different combinations of growth factors and inhibitors to generate blastoids, which in multiple aspects resemble real blastocyst stage embryos, although until now none have been able to generate viable animals. Most notably, two recent reports used D-EPSCs, either as a sole stem cell source (Belmonte group, B-blastoid)^35^ or in combination with TSCs (Zernicka-Goetz group, ZG-blastoid)^36^ to generate blastoids. Importantly, a large proportion of the blastoid cells generated with only D-EPSCs showed expression of genes previously associated with post-implantation stage lineages and not cells of the blastocyst. We therefore re-analyzed the scRNA-seq data provided in these reports and aligned them to our existing sampled preimplantation cells, along with an additional dataset containing cells up to E7.5^45^, to generate a developmental trajectory spanning fertilization to gastrulation (Figure 3A, S3A), resource data also available for visualization in SCope^56^. ZG-blastoids were generated by combining either D-EPSCs or ESCs with TSCs, using a slightly modified version of the protocol established by Rivron *et al*. for making blastoids^43^ with ESCs and TSCs. Corroborating previous findings, we found that both ESCs and D-EPSCs are able to give rise to similar cell types in these blastoids (Figure S3B, S3C): to cells resembling the E4.5 blastocyst EPI and to cells most similar to E4.5 PE or postimplantation parietal endoderm (Figure 3A, S3C), albeit reportedly D-EPSCs give rise to PE-like cells more efficiently than ESCs^36^. However, using the ZG-blastoid forming method the authors saw no detectable contribution of D-EPSCs towards the TE lineage and blastoid formation was not observed when TSCs were omitted^36^. B-blastoids, made from only D-EPSCs, also contained cells (B-blastoid EPI) that grouped with E4.5 blastocyst EPI cells and cells (B-blastoid PE) that grouped towards E4.5 blastocyst PE or postimplantation parietal endoderm cells (Figure 3A, S3D). However, only 6.7% of B-blastoid cells (B-blastoid-TE) clustered close to the TE lineage, between blastocyst TE cells and postimplantation ExE (Figure S3E). The remaining 60% of B-blastoid cells consisted of two intermediate clusters (B-blastoid-intermediate-1 and 2) that did not align with any blastocyst cells but instead resembled most closely certain postimplantation stage embryo cells. In an *in vitro* blastoid culture, these cells could be mistaken for TE cells as they express Cdx2, Krt8 and Krt18. However, they also co-expressed T (Brachyury) suggesting an embryonic or extra-embryonic mesoderm identity (Figure 3B). Indeed, B-blastoid-intermediate-1 cells showed the highest correlation with E5.5 EPI (r = .28, p < .001), E6.5-E7.5 EPI (r = .29, p < .001) and ExE mesoderm (r = .33, p < .001) and B-blastoid-intermediate-2 cells with ExE mesoderm (r = .67, p = .001) and other mesodermal lineages: mixed mesoderm (r = .64, p = .001), intermediate mesoderm (r = .62, p = .001), nascent mesoderm (r = .48, p = .001), as well as a strong resemblance to mesenchyme (r = .42, p = .001) (Figure 3C). Nevertheless, the small subpopulation of B-blastoid-TE cells resembling blastocyst TE is intriguing, and leaves the door open to the possibility that D-EPSCs can contribute to the trophoblast lineage under B-blastoid forming conditions. However, it should be noted that these cells expressed very low levels of conventional TE markers such as Cdx2, Elf5 and Gata3, while also exhibiting a higher similarity to ExE ectoderm (r = .74, p =.001), than genuine TE (r = .62, p = .001) (Figure 3B, 3C).

**Fig. 3:**
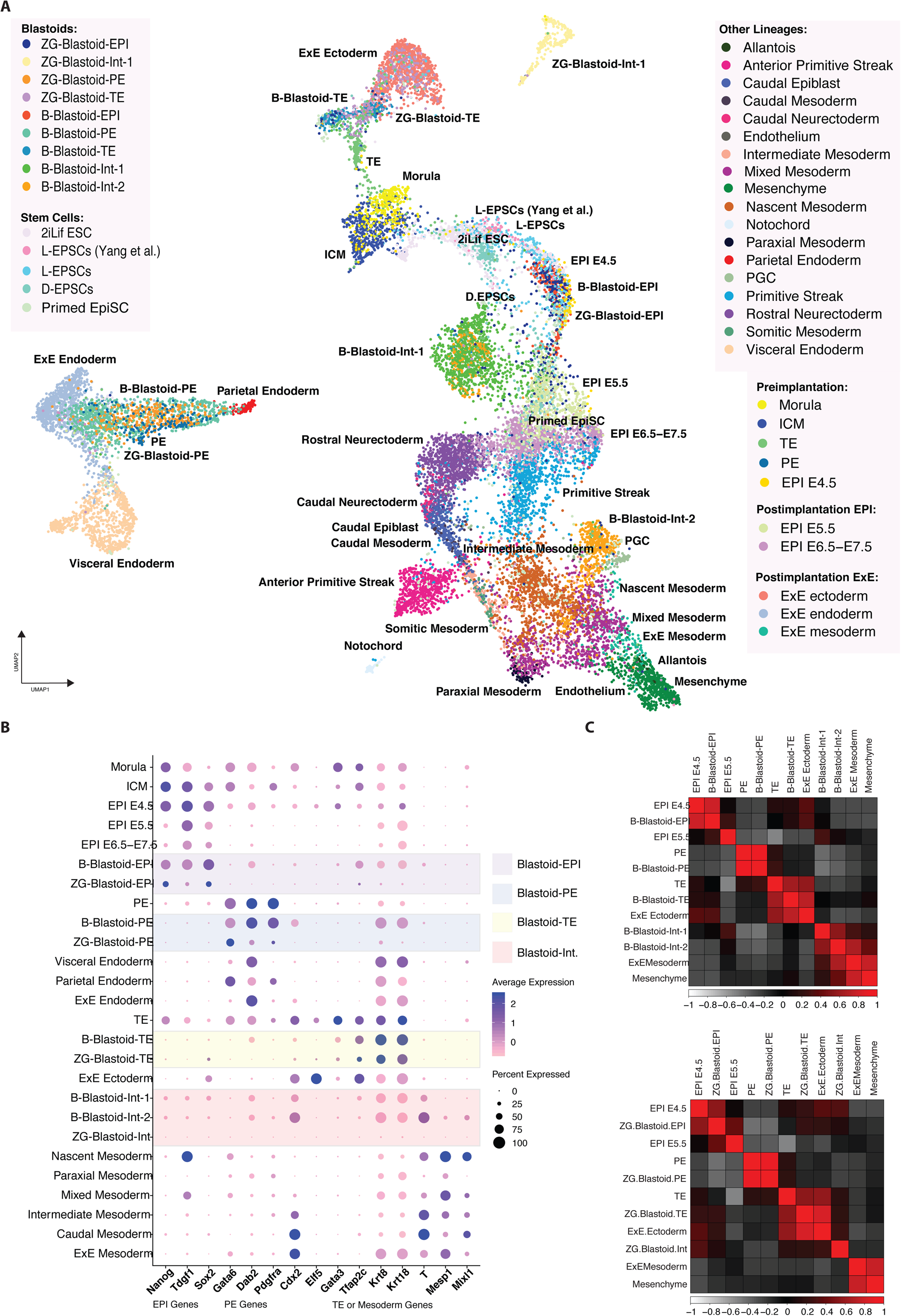
Single cell transcriptional comparison of blastoid cells to *in vivo* and different stem cell states. A) Single-cell UMAP tracking morula to E7.5 gastrulation in embryonic and extraembryonic lineages, comparing all stem cells, B-blastoid cells and ZG-blastoid cells. B) Dot plot representing frequency of expression and average expression for select lineage marker genes. C) Matrices showing correlation coefficients between B-blastoids (top) or ZG-blastoids (bottom) and selected lineages.

At the gene regulatory level, both B- and ZG-blastoid cells aligned well with embryo cells, indicating that the gene regulatory programs of natural embryos are recapitulated to a large extent in blastoids derived from D-EPSCs (Figure S4A). For example, the activity of selected regulons for lineage-specific TFs showed that blastoid-generated PE and EPI cells shared regulatory activity with their respective cell types in natural embryos (Figure S4B). Furthermore, in-depth analysis of target genes of NANOG and GATA4, EPI and PE TFs, respectively, showed similar target gene expression patterns between blastoids and natural embryos (Figure S4C). However, the TE regulatory state did not seem to be well recapitulated in D-EPSC-derived blastoids. Regulons associated with TE such as GATA3, CDX2, PITX1 and SOX6 are downregulated in blastoid TE compared to embryo TE (Figure S4B). Indeed, GATA3 target genes are downregulated in blastoid TE cells compared with natural embryos (Figure S4C), indicating that the misregulation of specific parts of the regulatory program underlying embryogenesis may limit blastoid development. We also investigated the regulatory activity of intermediate blastoid subpopulations to determine what may prevent these cells from becoming appropriate lineages present in the blastocyst. In line with gene expression analysis, we found that the intermediate blastoid populations (B-blastoid-intermediate-1 and 2) activated regulons of postimplantation EPI and mesodermal lineages (Figure S4A, S4B) such as T and MIXL1. We therefore analyzed gene expression of T target genes and found that B-blastoid-Intermediate-2 cells activate most, but not all, T targets (Figure S4C). At the same time, however, these cells, as well as mesodermal cells, also activate many targets of CDX2 (Figure S4C). This suggests that B-blastoid-intermediate-2 cells may have failed to activate the TE regulatory program and instead arrested between overlapping mesodermal and TE states. Altogether, these results demonstrate that the gene regulatory programs used in natural embryos are engaged to a large extent in EPSC-derived blastoids, but not fully, which might contribute to the developmental arrest of these structures.

### *In vivo* lineage contributions of totipotent blastomeres, L-EPSCs and D-EPSCs at embryonic day 4.5

The capacity to enter the trophoblast lineage can also be assessed *in vivo* by creating chimeras with a host embryo and analyzing lineage contributions at later developmental time points. To test lineage contributions of truly totipotent cells, we aggregated a morula-stage (E2.5) embryo (host) with a single blastomere from an 8-cell stage embryo (donor), as most, if not all blastomeres at the 8-cell stage are considered totipotent^46,47^. We allowed chimeras to develop for 48 hours before analyzing lineage contributions at the late blastocyst stage (E4.5). At E4.5 the three blastocyst lineages are clearly segregated (Figure 4A) and express well-characterized lineage specific markers, such as Sox2 (EPI), Sox17 (PE) and Cdx2, Gata3, Krt8, and Krt18 (TE)^48,49,50,51,16,52^. To visualize progeny of the donor cell, we isolated single blastomeres from embryos expressing either H2B-Gfp (nuclear-localized marker) or DsRed (no nuclear localization, marker appears in both cytoplasm and nucleus) and used wild-type embryos as hosts. We found that in 60-70% of chimeras the donor blastomere contributed to both the inner cell mass (EPI + PE) and the TE (Figure 4B) which was verified by co-immunostaining with the panel of lineage-specific markers (Figure 4C). These data serve as a benchmark for lineage contributions of truly totipotent cells in a chimera.

**Fig. 4:**
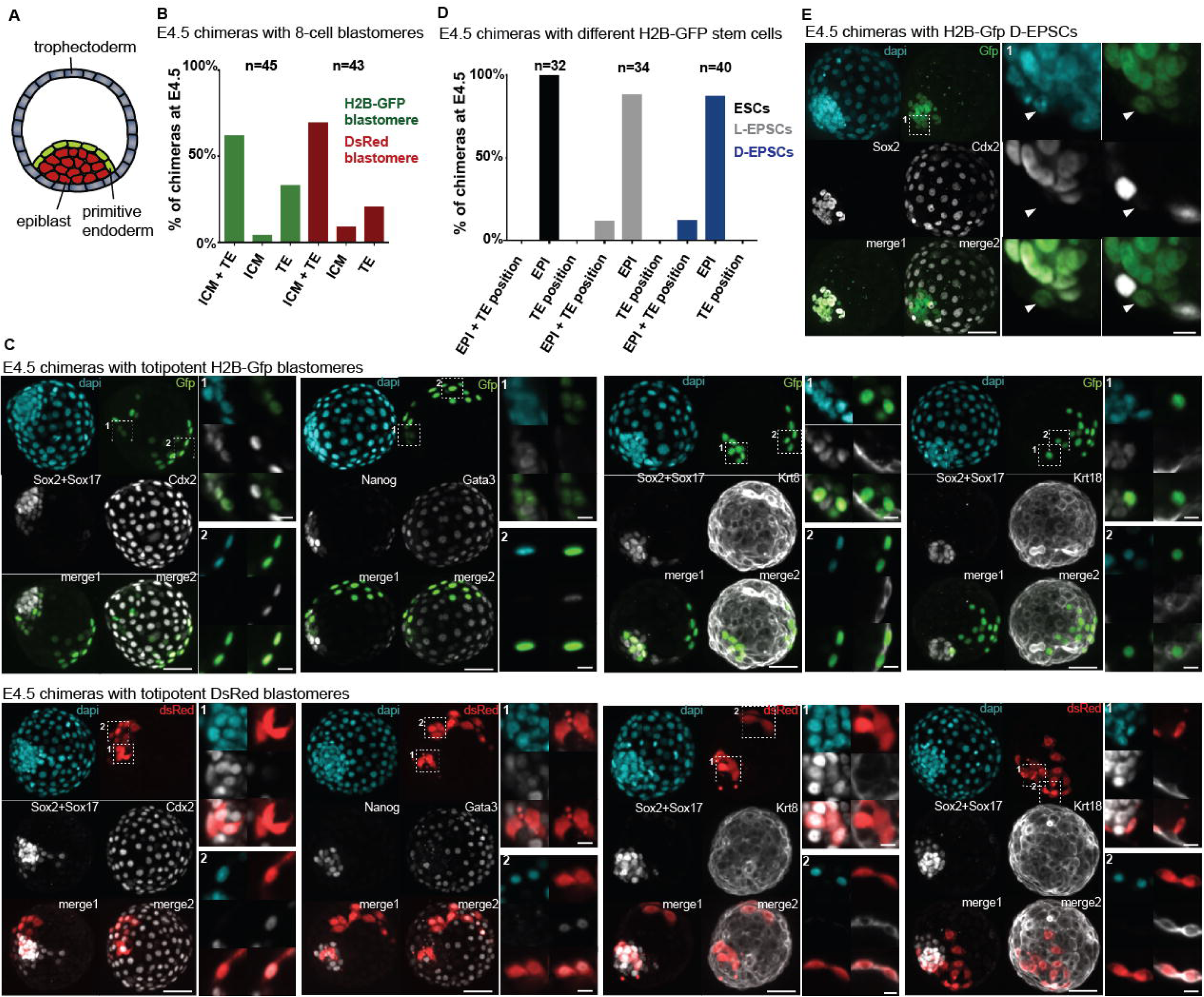
*In vivo* potential of candidate totipotent stem cells to give rise to the trophectoderm lineage at E4.5. A) Cartoon showing cell types of the E4.5 blastocyst embryo. B) Bar chart showing percent of E4.5 chimeric embryos with different lineage contributions. Chimeras were generated by aggregating a wild-type host embryo and a single blastomere of an 8-cell stage embryo that expressed either H2B-Gfp (green) or DsRed (red). n indicates number of chimeras generated. C) Immunofluorescent stainigs of E4.5 chimeric embryos with lineage contributions to both ICM and TE generated with a single blastomere of an 8-cell stage embryo that expressed either H2B-Gfp (top panel) or DsRed (lower panel). Stained for Sox2+Sox17 (ICM), Nanog (EPI), Cdx2, Gata3, Krt8 and Krt18 (TE) and Gfp or DsRed. Left panel shows maximum intensity projection of whole chimera, scale bar: 40 µm. Right panels show magnified single plane images of ICM and TE (bottom) contributions, scale bar: 10 µm. D) Bar chart showing percent of E4.5 chimeric embryos with different lineage contributions. Chimeras were generated by aggregating a wild-type host embryo with H2B-Gfp expressing ESCs (black), L-EPSCs (grey) or D-EPSCs (blue). n indicates number of chimeras generated. E) Example of E4.5 chimeric embryos generated with H2B-Gfp expressing L-EPSCs stained for Sox2 (EPI), Cdx2 (TE) and Gfp. Left panel shows maximum intensity projection of whole chimera, scale bar: 40 µm. Right panel shows magnified single plane image of Gfp positive L-EPSC contributions, scale bar: 10 µm.

We then aggregated L-EPSCs, D-EPSCs or control parental ESCs to wild-type host embryos and analyzed chimeras at E4.5. Interestingly, we observed that progeny of both L-EPSCs and D-EPSCs localized to trophectodermal positions in ∼20% of chimeras, while progeny of the parental ESC line cultured in 2iLif conditions localized only to the epiblast, corroborating previous studies^33,32^ (Figure 4D). However, when we immuno-stained chimeras for epiblast (Sox2) and trophectoderm markers (Cdx2), none of the L-EPSC or D-EPSC derived cells in the TE position showed expression of either marker (Figure 4E and S5). Therefore, we conclude that EPSCs can contribute cells that localize to the TE but do not express a key TE marker.

### *In vivo* lineage contributions of totipotent blastomeres, L-EPSCs and D-EPSCs at embryonic day 6.25

To confirm that the observed lineage contributions at E4.5 persist later in development, we examined chimeras post implantation. Shortly after implantation the EPI forms a cup-shaped epithelium, the PE forms the two layers of the visceral and parietal endoderm and the TE cells overlying the EPI proliferate to form the extraembryonic ectoderm (ExE) and the ectoplacental cone (Figure 5A). Before gastrulation is initiated at ∼E6.5, the boundaries of the different compartments are easily discernable, prompting us to analyze lineage contributions at E6.25. First, we generated chimeras with H2B-Gfp expressing blastomeres and showed that progeny of the blastomeres can contribute to both the Oct4-positive EPI, the Tfap2c and Elf5-positive ExE and the Tfap2c-positive ectoplacental cone (Figure 5B). Next, we generated chimeras with H2B-Gfp expressing L-EPSCs, D-EPSCs or ESCs and performed similar lineage analysis at E6.25. We found that all cell types readily contributed to the EPI lineage of the host embryo (Figure 5C). Interestingly, while ESC and D-EPSC chimeras occasionally contained donor cells in the trophoblast compartment (∼5% of chimeras), we also found that around 25% of L-EPSC chimeras contained a few cells in the ExE. However, when we performed immuno-staining for lineage-specific markers, cells localized to trophoblast regions did not express trophoblast markers such as Elf5 and Tfap2c (Figure 5D and S5F). Instead, most of these mis-localized cells expressed the EPI marker Oct4. These data emphasize that donor cell localization alone does not necessarily indicate appropriate lineage-specific marker allocation and therefore questions functional integration into the tissue. We also show an increased frequency of mis-localized L-EPSCs in chimeras, which may potentially explain the previously reported behavior of these cells.

**Fig. 5:**
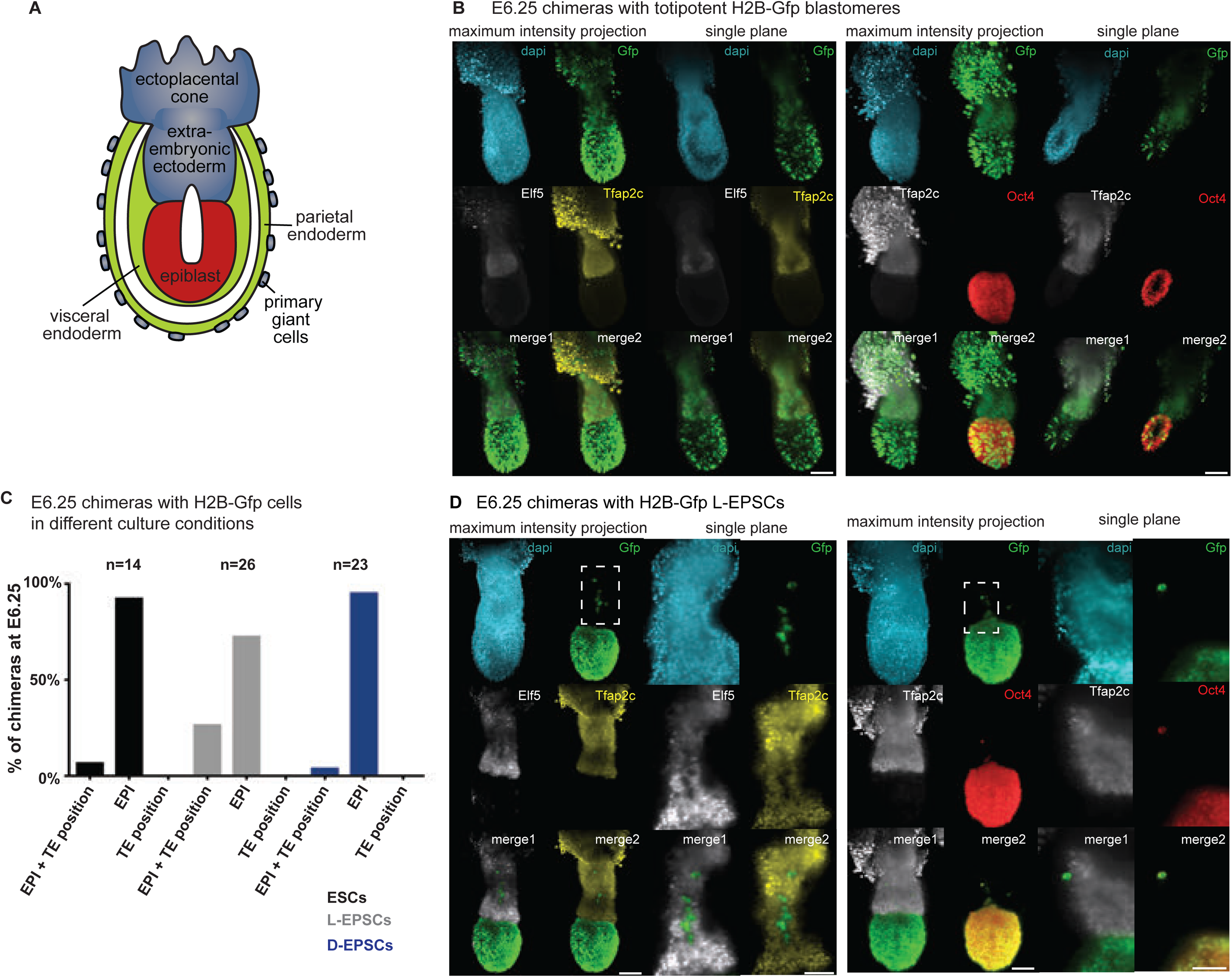
*In vivo* potential of candidate totipotent stem cells to give rise to the trophoblast lineage at E6.25. A) Cartoon showing cell types of the E6.25 embryo. B) Immunofluorescent stainigs of E6.25 chimeric embryos with lineage contributions to both EPI and trophoblast lineages generated with a single blastomere of an 8-cell stage embryo that expressed H2B-Gfp. Stained for Oct4 (EPI), Tfap2c and Elf5 (trophoblast compartment) and Gfp. Left panel shows maximum intensity projection of whole chimera, and right panels show single plane images, scale bar: 100 µm. C) Bar chart showing percent of E6.25 chimeric embryos with different lineage contributions. Chimeras were generated by aggregating a wild-type host embryo with H2B-Gfp expressing ESCs (black), L-EPSCs (grey) or D-EPSCs (blue). n indicates number of chimeras generated. D) Examples of E6.25 chimeric embryos generated with H2B-Gfp expressing L-EPSCs stained for Oct4 (EPI), Tfap2c and Elf5 (trophoblast compartment) and Gfp. Left panel shows maximum intensity projection of whole chimera, scale bar: 100 µm. Right panel shows magnified single plane image of Gfp positive L-EPSC contributions, scale bar: 50 µm.

### *In vivo* lineage contributions of totipotent blastomeres, L-EPSCs and ESCs in embryonic day 12.5 placentas

To test whether ExE-localized donor cells in chimeras give rise to differentiated trophoblast cell types, we analyzed chimeric placentas at E12.5. The placenta has a complex structure and contains both trophoblast as well as embryo-derived cell types^53^ (Figure 6A). Additionally, due to its high metabolite content, the placenta exhibits elevated levels of autofluorescence. These properties make immuno-fluorescent lineage analysis in the placenta a tricky task, requiring thorough evaluation aided by appropriate positive and negative controls. First, we identified antibodies and immuno-fluorescent staining conditions to label the different cell types of the placenta. We used Mct1 and Mct4 to label syncytiotrophoblast I and II, respectively, Tpbpa to label spongiotrophoblast, and Krt8, Cdh3 and Tfap2c to label all trophoblast cell types in both the spongio and the labyrinth zones. Tfap2c is a nuclear-localized TF, making it an ideal marker to detect co-localization with nuclear-localized lineage tracers (e.g. H2B-Gfp). Finally, we used CD31 to label embryo-derived endothelial cells in the placenta. We then used this marker panel to show that in chimeric placentas generated with a single H2B-Gfp or DsRed expressing totipotent blastomere and a wild-type host embryo, blastomere progeny contribute to all analyzed lineages (Figure S6). To unambiguously distinguish between trophoblast and embryo-derived cells in the placenta we took advantage of a technique termed tetraploid complementation, which involves generating a chimera using a tetraploid host embryo and diploid ESCs^54,55^ (Figure 6B). Tetraploid cells are not tolerated in the embryonic compartment and ESCs do not contribute to the trophoblast compartment. Therefore, any surviving conceptus at E12.5 consists of trophoblast originating from tetraploid cells and embryonic tissues originating from ESCs. We generated chimeras in which either the tetraploid cells or ESCs (Figure 6C) carried H2B-eGfp and immuno-stained placental sections at E12.5 for the markers described above (Figure S7). As expected, we saw that in chimeras with H2B-eGfp-posititve tetraploid cells trophoblast markers (Tfap2c, Cdh3, Tpbpa, and Mct4) always overlapped with the Gfp signal, while the embryonic marker CD31 did not. In contrast, in chimeras with H2B-eGfp-posititve ESCs only CD31 overlapped with the Gfp signal, and trophoblast markers were excluded from Gfp-positive cells. This panel highlights the difficulty in distinguishing different cell types in the placenta, especially in the labyrinth zone, without detailed analysis of markers and also emphasizes the challenge of matching a nuclear label with a membrane-localized signal in individual cells. Co-localization can be interpreted more clearly when the fluorescent lineage tracer and the cell-type specific marker are in the same sub-cellular compartment, as exemplified in our staining panel by the co-localization of H2B-eGfp and Tfap2c. Next we generated chimeric placentas using diploid host embryos and L-EPSCs and analyzed them using the same marker panel (Figure 6D). We could only detect Gfp-positive cells in the embryonic, but not in the trophoblast compartment, suggesting that L-EPSCs do not readily give rise to differentiated trophoblast cell types.

**Fig. 6:**
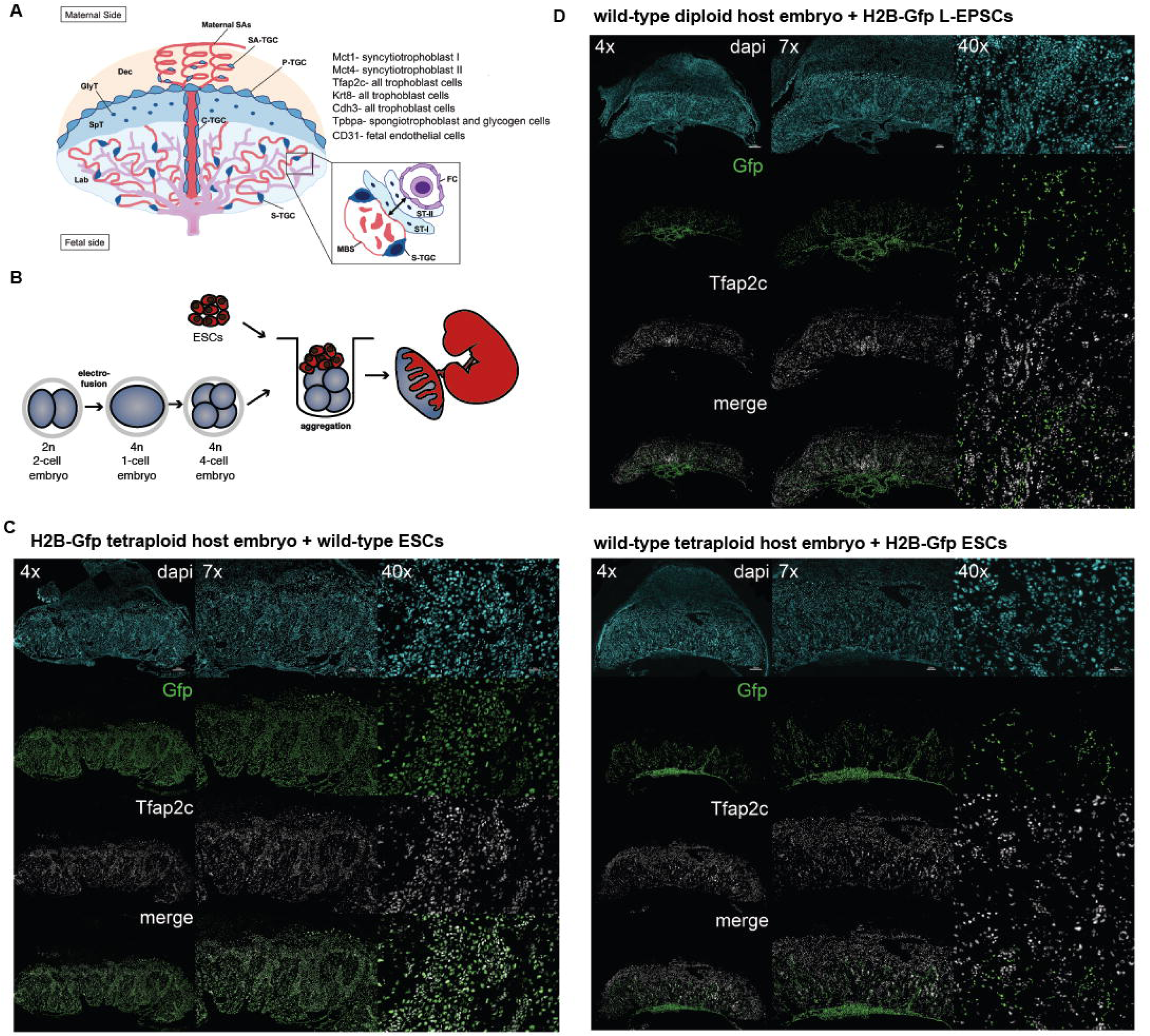
*In vivo* potential of candidate totipotent stem cells to give rise to the trophoblast lineage of the E12.5 placenta. A) Cartoon indicating cell types and organization of the E12.5 placenta. Markers of different cell types used in this study are indicated. B) Experimental design of tetraploid complementation assay. Tetraploid host embryos are generated by electro-fusing embryos at the 2-cell stage. A small clump of ESCs are aggregated with tetraploid host embryos to generate a chimera. Chimeric placentas are analyzed at E12.5. C) Examples of chimeric E12.5 placentas generated with H2B-Gfp expressing tetraploid host embryo and wild-type ESCs (left panel) and wild-type tetraploid host embryo and H2B-Gfp expressing ESCs (right panel). Placenta sections are immuno-stained for Gfp and Tfap2c, which labels all trophoblast cell types. Images shown at different magnifications. Scale bars: 500 µm (4x), 200 µm (7x), 50 µm (40x). D) Example of chimeric E12.5 placentas generated with wild-type host embryo (diploid) and H2B-Gfp expressing L-EPSCs. Placenta sections are immuno-stained for Gfp and Tfap2c. Images shown at different magnifications. Scale bars: 500 µm (4x), 200 µm (7x), 50 µm (40x).

## Discussion

Here we present different criteria to evaluate the differentiation potential of early embryonic stem cells. We provide a large compiled dataset of single cell transcriptomes covering different *in vivo* cell types from fertilization to gastrulation, as well as several early stem cell types, which can be used to map a novel cell type based on transcriptional similarities. As a resource for the community, the data presented here is made available through a user-friendly file format that can be explored using the single-cell analysis tool SCope^56^. Users can upload the .loom files provided here (https://github.com/pasquelab/totipotency) enabling to browse the data sets at will.

We demonstrate assays to directly test the differentiation potential of cells by converting or reprogramming cells in vitro to TSCs or by analyzing lineage contributions to the trophoblast compartment *in vivo* in the context of a chimera. Of note, in the later experiments we only focused on extraembryonic contributions to the trophoblast lineage and therefore cannot draw conclusions about potential contributions to the extraembryonic endoderm lineage. By using totipotent blastomeres as examples of a truly totipotent state, we highlight the importance of evaluating chimeric contributions not only based on the localization of donor cells, but also on the expression on lineage-specific marker analysis.

Using these criteria, we examine the potential of two novel stem cell states (L-EPSCs and D-EPSCs) that have been reported to have expanded/extended potential beyond pluripotency. Surprisingly, we fail to find convincing evidence that these cell types harbor extensive expanded or extended potential. Instead, based on our transcriptomic comparison L-EPSCs most closely resemble E4.5 EPI cells or the parental ESCs cultured in 2iLif and D-EPSCs the E5.5 EPI or EpiSCs. In TSC conversion or reprogramming settings neither L-EPSCs or D-EPSCs show enhanced potential compared to parental ESCs. Finally, in chimeric experiments L-EPSCs and D-EPSCs only show convincing contribution to the EPI lineage. Interestingly, we found that in chimeras analyzed at the late blastocyst stage generated with both L-EPSCs or D-EPSCs, but not ESCs, cells occasionally localized to TE positions but did not express either EPI nor TE markers. We hypothesize that these mis-localized, marker-negative cells are not maintained long term and are likely in the process of getting eliminated from the compartment where they do not belong. We also detected mis-localized cells in chimeras made with L-EPSCs just prior to the onset of gastrulation. Majority of these cells however continued to express Oct4 and lacked trophoblast marker expression. Therefore, it is likely that these cells are not the progeny of the TE-localized, marker-negative cells observed in blastocyst-stage chimeras. Instead, mis-localization of L-EPSCs may occur during postimplantation development and coud potentially be due to weak anchorage or un-synced developmental timing, allowing spurious integration.

Our results which suggest that L- and D-EPSCs are unable to enter the trophectoderm lineage are seemingly in contrast with the recent study reporting the formation of blastocyst-like structures, termed blastoids, using only D-EPSCs^35^ or L-EPSCs. We therefore carefully re-examined the cell types generated in blastoids. Our results corroborate the idea that EPSCs are able engage the gene regulatory programs utilized in distinct cellular lineages in natural embryos. However, we also found differences between the gene regulatory programs of natural embryos and EPSC-derived blastoids, which were most apparent in the TE lineage. Our re-analysis of the scRNA-seq data of B-blastoids made form D-EPSCs indicated that only 6.7 % of cells sequenced were categorized as TE and even these showed an ExE-like profile. These TE-like cells also failed to show robust expression of classical TE markers such as Cdx2, Gata3 or Elf5. Problematically, the most abundant cell types in B-blastoids (B-blastoid intermediate 1 and 2) seem to most closely resemble mesoderm, expressing markers such as T, yet also share a number of common markers with the trophoblast lineage, such as Cdx2, Krt8, Krt18, Tfap2c. With such cell composition it is not surprising that B-blastoids are not able to generate a live conceptus. Although not abundant, the presence of TE or ExE-like cells in B-blastoids is intriguing and leaves the door open for the possibility that some EPSCs may indeed harbor potential to differentiate into trophoblast. Of note, ESCs are also able, in some cases, to form blastoids^44,36^ but whether they also recapitulate the gene regulatory programs of natural embryos like EPSC-derived blastoids do remains to be determined.

Why is this potential only revealed in the blastoid-forming assays and not in the context of chimeras? Forcing cells to the surface of a forming sphere in the blastoid method may mimic TE-inducing cues better than aggregation assays, which allow positional freedom of aggregated cells within the host embryo^3^. Positional freedom permits cells to group with TE or ICM compartments in the forming blastocyst based on their identity, therefore if only the potential exists for TE fate, this may not be realized in an aggregation setting. Additionally, B-blastoids are formed under specific culture conditions which may direct differentiation more robustly than the environment of the embryo. Supporting this notion, D-EPSCs were not able to give rise to a TE-like layer under a different blastoid forming protocol (ZG-blastoid). Instead, TSCs had to be used^36^. It should however still be considered that the TE or ExE-like blastoid cells fail to express Cdx2, Gata3 and Elf5 transcripts in similar levels to endogenous TE or ExE of the embryo suggesting that their transcriptional profile still is distinct from *in vivo* cells.

Notably, the B-blastoid method employs Bmp4 and inhibits Activin/Nodal signaling^35^, conditions which were also used in another blastoid protocol by the Tomoda lab^44^. The Tomoda group used EpiSCs as starting cells and were also able to produce blastoids with certain TE-like marker expression in the surface layer. Additionally, high Bmp4 and low Activin/Nodal was previously computationally predicted and shown *in vitro* to activate TE-marker gene expression in ESCs in which Jak/Stat signaling was inhibited^57^. These data suggest that high Bmp4 and low Activin/Nodal signaling may be key to TE-like cell induction.

Importantly, these signaling conditions are also involved in inducing proximal mesoderm fates during gastrulation and Bmp induces mixed mesoderm and trophoblast differentiation in EpiSCs and hESCs^58^, consistent with the appearance of abundant mesoderm-like cells in B-blastoids. Could the starting stem cell state be crucial for facilitating mesoderm versus trophoblast differentiation? Indeed, it was shown that Cdx2 overexpression in ESCs induces reprogramming into TSCs, while Cdx2 overexpression in EpiSCs results in mesodermal gene expression^19,59^, highlighting the importance of the starting state for different differentiation outcomes. Notably, as our analysis placed D-EPSCs closest to primed EpiSCs, the widespread induction of mesodermal profiles is not surprising. We propose that to truly unlock a cell’s differentiation potential into any extraembryonic or embryonic lineage, a starting state more resembling earlier embryonic, such as morula stages is needed. Our study highlights this challenge and sets gold standards for evaluating the differentiation potential of cells using various methods.

## Methods

### ESC and EPSC culture

Naïve mouse ESCs and D-EPSCs were cultured in a base medium of N2B27 prepared as follows: 1:1 ratio of DMEM/F12 (ThermoFisher Scientific 21331020) and Neurobasal (ThermoFisher Scientific 21103049); 1 mL N2 supplement (ThermoFisher Scientific 17502001) or 1xLNDiff Neuro2 supplement (Gibco, 17502048); 2 mL B27 supplement minus vitamin A (ThermoFisher Scientific 12587-010 or 17504044); 1x Glutamax (ThermoFisher Scientific 35050061); 0.1mM β-mercaptoethanol (ThermoFisher Scientific 21985023). For best results, this base media was further supplemented with 1-5% knockout serum replacement (KSR; ThermoFisher Scientific 10828028) as described^60,33^. Both naïve ESCs and D-EPSCs were cultured at 20% O2 and 5% CO2 at 37°C on mitomycin C (MMC; Sigma-Aldrich M0503) inactivated mouse embryonic fibroblast (MEF) feeder cells (approx 30,000 cells/cm2). E12.5 DR4 MEFs were routinely plated on 0.2% gelatin-coated plates in ESC base media prepared as follows: DMEM (ThermoFisher Scientific 11960069); 15% foetal bovine serum (FBS; Wiscent); 1x Glutamax (ThermoFisher Scientific 35050061); 1x non-essential amino acids (NEAA; ThermoFisher Scientific 11140-050); 1 mM sodium pyruvate (ThermoFisher Scientific 11360070); 0.1 mM β-mercaptoethanol (ThermoFisher Scientific 21985023 or Merck, M3148 SIGMA). MEF plates were used within 1 week and washed with DPBS (ThermoFisher Scientific 14190250) prior to plating of ESC/D-EPSCs in appropriate media.

For naïve ESC culture, N2B27 (1-5% KSR) base media was supplemented with 1 µM PD0325901 (Tocris 4192); 3 µM CHIR99021 (Tocris 4423); and 1000 U/mL mouse LIF (generated in-house). For D-EPSC culture, N2B27 1-5% KSR base media was supplemented with 1x NEAA (ThermoFisher Scientific 11140-050); 10 ng/mL recombinant human LIF (hLIF; Peprotech 300-05); 3 µM CHIR99021 (Tocris 4423); 2 µM Dimethindene maleate (Tocris #1425); and 2 µM Minocycline Hydrochloride (Santa Cruz #sc-203339). Media was changed daily for both ESCs and D-EPSCs, with single-cell passaging every 2-3 days using accutase (ThermoFisher Scientific A1110501) at split ratios between 1:5 and 1:12.

L-EPSCs were cultured in a base media prepared as follows: DMEM/F12 (ThermoFisher Scientific 21331020); 20% KSR (ThermoFisher Scientific 10828028); 1× Glutamax (ThermoFisher Scientific 35050061) (or DMEM/F12 (Gibco, 13320074), 20%LKnockOut Serum Replacement (KSR, Gibco, 10828028), 2.25 mM L-glutamine); 1× NEAA (ThermoFisher Scientific 11140-050); 0.1 mM β-mercaptoethanol (ThermoFisher Scientific 21985023 or Merck, M3148 SIGMA). This base media was supplemented with 10 ng/mL hLIF (Peprotech 300-05) or 1000 U/ml homemade mouse Lif; 1 µM PD0325901 (Tocris 4192 or Axon 1408); 3 µM CHIR99021 (Tocris 4423, or Axon 1386); 4 µM TCS JNK 60 (Tocris 3222); 5 µM XAV939 (Sigma-Aldrich X3004); 10 µM SB203580 (Tocris 1402); and 0.3 µM A-419259 (Santa Cruz sc-36109; or Tocris 39142). L-EPSCs were cultured at 20% O2 and 5% CO2 at 37°C on MMC-inactivated SNL76/7 feeder cells or MMC-inactivated MEF feeders (∼50-80,000 cells/cm^2^). SNL feeders were plated on 0.2% gelatin-coated plates in ES base media (described above). SNL plates were used within 3-4 days and washed with DPBS prior to plating L-EPSCs. Media was changed daily, with single-cell passaging every 3 days using accutase at split ratios between 1:3 and 1:12.

### Conversion to EPSC conditions

To convert naïve ESCs into either D-EPSC or L-EPSC conditions, cells were passaged and plated at range of low cell densities (1-3,000 cells/cm2) on either MMC-inactivated E12.5 WT or DR4 MEFs or SNL feeders in their original media. The following day the media was changed to D-EPSC or L-EPSC conditions, respectively. The cells were passaged when approaching confluence (4-5 days) at a range of split ratios (e.g. 1:3, 1:5, 1:8). D-EPSCs and L-EPSCs were cultured for at least 5 passages (∼2 weeks) prior to use in chimaera or *in vitro* potency experiments.

### TSC culture

TSCs were cultured in standard conditions: RPMI-1640 (Sigma-Aldrich R0883); 20% FBS (Wiscent); 1x Glutamax (ThermoFisher Scientific 35050061); 1 mM sodium pyruvate (ThermoFisher Scientific 11360070); 0.1 mM β-mercaptoethanol (ThermoFisher Scientific 21985023). This TS base media was supplemented with 25 ng/mL FGF4 (R&D Systems 235-F4) and 1 µg/ml heparin (Sigma-Aldrich H3149). TSCs were routinely cultured on MEFs plated on 0.2% gelatin-coated plates (approx 30,000 cells/cm^2^). Medium was changed every 1-2 days and cells passaged before reaching confluency (every 3-4 days) at split ratios between 1:5 and 1:12.

### Cell line generation

H2B-eGfp ESCs (ICR) were derived directly from E3.5 blastocysts (JAX 006069 backcrossed to ICR in-house^61^). Briefly, single E3.5 blastocysts were transferred into a well containing 100 µL 2iLif media (96-well plate pre-coated with E12.5 DR4 MEFs). Embryos were left undisturbed for approx 48 hours at which point additional 100 µL 2iLif media was added per well. The top 100 µL of media was then changed every two days until the outgrowth reached passaging size (∼5-7 days). Outgrowths were passaged using 50 µL accutase per well and plated into a new 96-well MEF plate. Cells were then passaged as described above until expanded for morphological assessment and cryopreservation of selected clones.

R1-mScarlet-NLS ESCs were generated from R1-ESCs (129X1 × 129S1;^62^) by transfection (Lipofectamine 2000; ThermoFisher Scientific 11668027) with PiggyBac CAG-mScarlet-NLS construct and isolation of single clones.

Wild-type TSCs were TS-F4 (ICR) described previously^63^. Elf5-2A-mCherry TSCs were generated by transfection (JetPRIME; Polyplus 114-07) of TS-F4 cells with vector expressing Cas9 and gRNA targeting proximal to Elf5 stop codon. Donor construct containing desired insert (GSG-P2A-SV40-NLS) and selection cassette (SV40::NeoR) flanked by homology arms was co-transfected. Cells were selected with 100-200 µg/mL G418 (ThermoFisher Scientific 10131027) and correctly targeted clones identified by PCR genotyping and Sanger sequencing. Floxed selection cassette was subsequently removed by transfection with Cre and desired clones identified by PCR genotyping and Sanger sequencing. Heterozygous Elf5-2A-mCherry TSC clones were used in this study and faithful overlap between Elf5 and mCherry protein validated by immunostaining.

5ECER4G20 eGfp-Cdx2-ER (iCdx2) ESCs were a kind gift from H. Niwa^19^. These cells constitutively express eGfp and are heterozygous for *Oct4* (*Pou5F1*). Heterozygous Elf5-2A-mCherry iCdx2 ESCs were generated as above, except for use of SV40::HygroR selection cassette and Lipofectamine based transfection.

### ESC-to-TSC reprogramming

For *in vitro* testing of EPSC potency, iCdx2 Elf5-2A-mCherry ESCs were converted to either D-EPSC or L-EPSC conditions as described above. To initiate ESC-to-TSC reprogramming, 50-200 cells/cm^2^ were plated onto low density E12.5 DR4 MEFs (∼10,000 cells/cm^2^ on gelatin-coated plate) in their original media. The following day the media was changed to TSC media (+Fgf4/Heparin) with or without 1 µg/mL 4-hydroxytamoxifen (Sigma-Aldrich H7904) to induce Cdx2. Media was changed daily and the degree of TSC reprogramming assessed by flow cytometry on day 6.

For ESC-to-TSCs differentiation followed by RT-qPCR, L-EPSCs and ESCs were gradually feeder-depleted by passaging every two days with 0.1% gelatin (porcine skin, 0.1%□g/v final, Sigma, G2500) and a feeder percentage of 100%, 75%, 50% to 0% at every passage in L-EPSCM and 2iLif, respectively. After complete feeder removal, the cells were cultured at a density of 1×10^5^□cells on gelatin-coated culture 6 well plates in TSC medium (as above). The cell culture medium was refreshed every two days and cells were collected every three days from D0 to D12 for RT-qPCR analysis of TSC marker genes expression using the following primers:

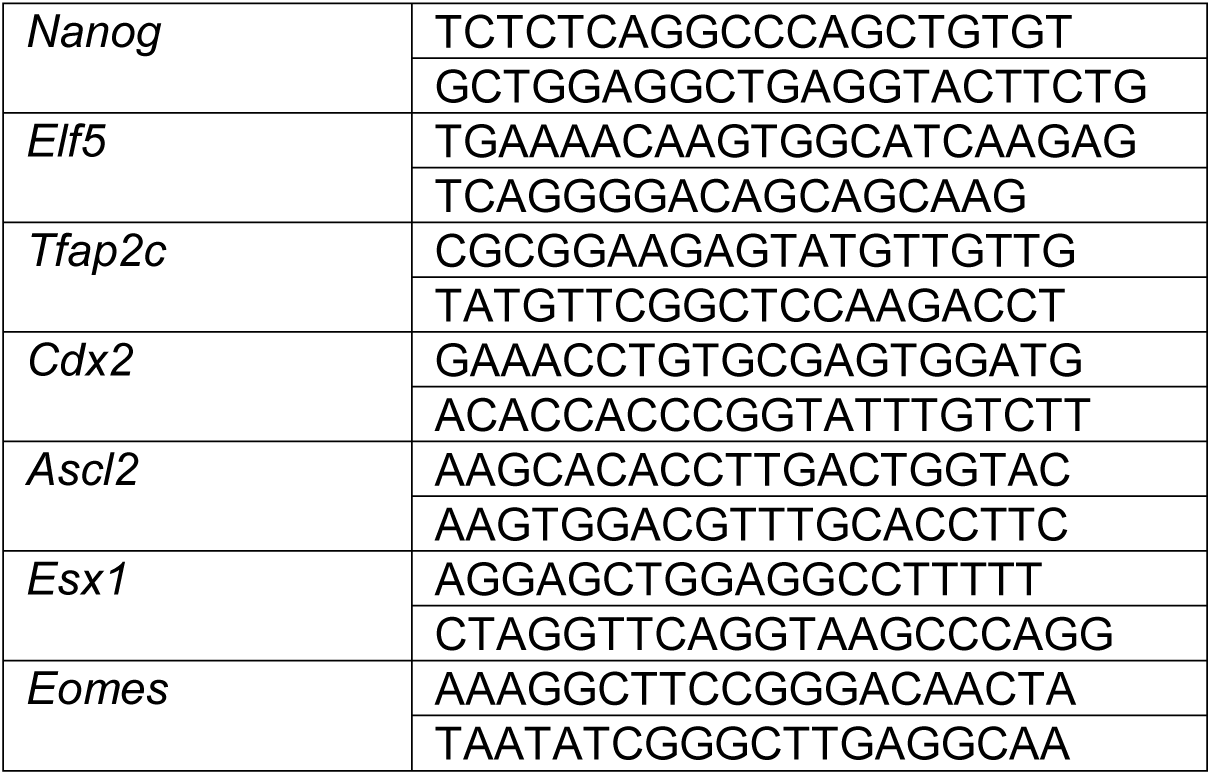

### Flow cytometry

Flow cytometry assay was designed to assess differential protein levels of TSC surface markers CD40^63^ and Plet1^42^, as well as TSC transcription factor Elf5^20^ using a reporter cell line. Titration experiments were carried out to optimize staining against CD40 (1:20; BD Biosciences 740700) as well as Plet1 (1:200; Nordic MUbio MUB1512P conjugated to Bio-Rad ReadiLink 633/655 1351005). Single-cell suspensions for ESCs and EPSCs undergoing ES-to-TSC reprogramming as well as control cell lines (ESC/EPSC day 0 samples; wild-type TSCs and Elf5-2A-mCherry TSCs) were generated using 0.25% trypsin (ThermoFisher Scientific 25200072). Cells were incubated with conjugated antibody mix in PBS/2% FBS for 60 min on ice. Post-staining, cells were washed twice with PBS/2% FBS prior to resuspension in PBS/2% FBS supplemented with 0.5 µm Sytox blue (ThermoFisher Scientific S11348) to assess cell viability. Cells were transferred to flow tubes with 40 µm cell strainer lid and analyzed using BD LSR II at the SickKids Flow Cytometry facility. For ES-to-TSC reprogramming samples, at least 100,000 events were recorded and analyzed using FlowJo software. Gates were drawn based on fluorescence minus one (FMO) controls.

### Bulk RNA sequencing library preparation

Total RNA was isolated from 2iLif ESCs and L-EPSCs reprogramming intermediates at indicated days using TRIzol reagent according to the manufacturer’s protocol. 4 µg of total RNA was used to generate libraries using of stranded poly(A) mRNA-Seq library with the KAPA stranded mRNA Library prep kit (KAPA Biosystems, KK8421). Libraries concentration was quantified with Qubit dsDNA HS (High Sensitivity) Assay Kit (Invitrogen, Q32854), and the final pool was generated by combining individual libraries in equimolar ratio. Libraries were sequenced on an Illumina HiSeq 4000 instrument (Illumina) with 51 bp reads and read depth ranging ∼33 mln reads. Sequencing reads were mapped to mm10 reference genome using STAR 2.5.3a^64^. On average, 77,16% of reads were uniquely mapped and only those were kept for further analyses. Subsequently, the featureCounts function from the R Bioconductor package “Rsubread” (version 1.5.2)^65^ was used to assign mapped reads to genomic features.

### Bulk RNA sequencing analysis

Processing raw read counts was performed as described in^66^. Briefly, the DESeq2 package and the associated protocol^67^ was used. Only genes that express at least 10 reads in total across all libraries were retained. PCA was performed using plotPCA function from the DESeq2 package with input of top 500 most variable genes after rlog transformation. Unless mentioned otherwise, gene expression was presented as log2 values after size-factor normalization for the differences in library size (DESeq2).

### Integration of bulk RNA-seq data with published single-cell RNA-seq data

Principal component analysis of top 500 most variable genes. Single-cell data from published datasets (Deng *et al*. 2014^37^; Posfai *et al*. 2017^3^; Yang *et al*. 2017^33^) were processed together with samples from this study. The reads from single-cell samples were averaged within the same embryonic timepoint. Subsequently, all samples were normalized together for the library size using size-factor normalization in the DESeq2 package.

Comparison of gene expression of different L-EPSC conversion timepoints to 4, 8 and 16 cell stage preimplantation embryos was done in two-steps. First, stage-specific markers were defined using k-means clustering (SC3) of single-cell data, followed by differential expression analysis between the clusters. Second, these markers were used for the comparison between L-EPSC timepoints and the corresponding embryo stages in the averaged single-cell data.

### Library preparation for single-cell RNA sequencing

Single-cells from stem cells were sorted by FACS into 384 well plates containing lysis and RT, whereas embryo cells were manually picked and directly dispensed into lysis buffer containing RT. The current study generated cDNA libraries using the Smart-seq2 protocol, as previously described^68,69^.

### Single-cell RNA sequencing data pre-processing and quality control

Smart-Seq2 read files (E2.5 and E4.5 embryos, and three pluripotent stem cell conditions: ESCs cultured in 2iLif, L-EPSCs, and D-EPSCs) were mapped to the mouse reference genome (mm10) using STAR aligner^64^ and only uniquely mapped reads were used for expression level estimation as reads per kilobase of gene model and million mappable reads (RPKMs) using RefSeq annotation and previously established pipeline^70^. Cells were quality-filtered with minimum cut-off of 500 expressed genes per cell and Spearman’s correlation greater than 60% between cells. Pre-processed gene expression matrices were downloaded as provided by Deng *et al*., 2014^37^ (GSE45719), Posfai *et al*., 2017^3^ (GSE84892), Mohammed *et al*., 2017^38^ (GSE100597), Chen *et al*., 2016^39^ (GSE74155), Li *et al*., 2019^35^ (GSE135289, GSE135701), Yang *et al*., 2017^32^ (ERP005641), Pijuan-Sala *et al*., 2019^45^ (https://github.com/MarioniLab/EmbryoTimecourse2018), Sozen *et al*., 2019^36^ (GSE134240).

### Single-cell gene expression analysis of merged datasets

Analysis of the filtered data was conducted in R version 3.6.1 using Seurat suite version 3.1.0.^71^. Integration of single-cell data was performed using Seurat’s canonical correlation analysis (CCA) integration tool; datasets were scaled and log-transformed before selection of 2000 most variable genes which were used to compute principal component analysis (PCA). Manifold approximation and projection (UMAP) coordinates were calculated using the top 14 PCs. Seurat’s FeaturePlot function was used to demonstrate individual gene expression on UMAP embedding. Merged datasets were clustered, annotated, then downsampled prior to CCA integration. Correlation matrixes were constructed using corrplot v0.84. Clustering of preimplantation lineages was performed using Seurat’s shared nearest neighbor (SNN) algorithm implemented in FindClusters function. Differential expression analysis was performed using Wilcoxon’s rank sum test using a minimum cutoff of 0.45 average log fold change.

### Gene Regulatory Network inference

Gene regulatory networks were inferred using pySCENIC (0.9.15; python implementation of SCENIC)^40^ in Python version 3.6.9. First, raw expression data was normalized by dividing feature counts of each cell by the total counts for that cell and multiplying by factor of 10000 followed by log1p transformation. Subsequently, normalized counts were used to generate co-expression modules using GRBboost2 algorithm^72^ implemented in arboreto package (v0.1.3)^73^. Next, gene regulatory networks were inferred using pySCENIC (with default parameters and mm10 refseq-r80 10kb_up_and_down_tss.mc9nr and mm10 refseq-r80 500bp_up_and_100bp_down_tss.mc9nr motif collections) resulting in the matrix of AUCell values that represent the activity of each regulon in each cell. The resulting gene regulatory networks contained 451 regulons for merged datasets in Figure 1, S2 and S3 and 333 regulons for datasets in Figure 3, S3 and S4 and have been added to the .loom files which can be downloaded from: https://github.com/pasquelab/totipotency and browsed interactively by uploading the loom file on the SCope platform, www.scope.aertslab.org. The loom files contain non-integrated gene expression and regulon data.

For generating UMAP plots based on gene regulatory information in Figures S2E and S4A, the AUCell matrix was split by dataset of origin and integrated using Seurat’s canonical correlation analysis (CCA) integration tool. Anchors for integration were found using FindIntegrationAnchors function with default parameters and dims = 1:20 and data was integrated across all features. SCENIC-based UMAP were constructed using runUMAP function with default parameters except for dims = 25 and min.dist = 0.35 in Figure S2E and dims = 25 and min.dist = 0.25 in Figure S4A.

The list of target genes were downloaded from the loom file through the SCope platform.

### Mouse lines and embryos

ICR (breeding stock from Charles River, Montreal, Canada), *DsRed*^74^, *H2B-Gfp*^61^ mouse lines were used in this study. Embryos were collected at appropriate time points from 5-8 week old hormone-primed (5 IU each, pregnant mare serum gonadotropin (Sigma) and human chorionic gonadotropin (Sigma), 48 hours apart) and mated females. If not immediately used, embryos were cultured in small drops of KSOM supplemented with amino acids (EMD Milipore) under mineral oil (Zenith Biotech, Guilford, CT) at 37°C, with 5% CO2 for specified times. All animal work was carried out following Canadian Council on Animal Care Guidelines for Use of Animals in Research and Laboratory Animal Care under protocols approved by The Centre for Phenogenomics Animal Care Committee (20-0026H).

### Generation, culture and isolation of chimeras using diploid or tetraploid hosts

To isolate 8-cell stage blastomeres embryos from *H2B-Gfp* or *DsRed* mouse lines were harvested on the morning of day E2.5 by flushing the oviduct. The zona pellucida was removed using acid Tyrode’s solution (Sigma, Oakville, Canada) and embryos were washed in M2. Dissociation was performed by incubating embryos in TrypLE Select (Gibco™, Thermo Fisher Scientific, Waltham, MA) for 3-6 minutes at 37°C followed by pipetting through fine pulled glass capillaries. Individual cells were picked, washed in warm M2 and used as donor cells. For host embryos, E2.5 embryos were isolated and zona was removed in a similar way. To generate aggregation chimeras a single donor cell and a single host embryo were then brought together in a micro-well generated by pressing a blunt end needle into the bottom of a plastic tissue culture dish (Falcon™, Thermo Fisher Scientific) in drops of KSOM under oil. Such aggregation chimeras were either cultured for 48 hours (E4.5) or transferred into oviducts of pseudopregnant females on the following day and isolated either on day E6.25 or E12.5.

For generating chimeras with different stem cells (ESC, L-EPSC, D-EPSC), host embryos were generated as described above. A small clump (6-8 cells) of stem cells expressing H2B-Gfp were aggregated to each host embryo in a micro-well. Stem cell clumps were made by briefly trypsinizing the cells, inactivating or diluting trypsin with media and manually picking appropriate-sized clumps using a fine glass capillary. Chimeras were cultured or transferred into pseudopregnant females as above.

For tetraploid complementation experiments embryos from either wild-type or H2B-Gfp mouse lines were isolated at the 2-cell stage (E1.5). Cells were electro-fused into one tetraploid cell using a Cell-fusion instrument, CF-150B pulse generator with 250 μm electrode chamber (BLS Ltd, Hungary). The procedure is described in detail in^75^. Such tetraploid embryos were cultured for one day before using them as host embryos. In tetraploid complementation experiments a small clump (6-8 cells) of ESCs either from a wild-type line or an H2B-Gfp expressing cell line was used. Aggregation, chimera culture and transfer were performed as before.

### Immunofluorescent staining of E4.5 and E6.25 chimeras

Whole mount immunofluorescence staining of E4.5 embryos was performed as previously described^3^. Briefly, embryos were fixed in 4% paraformaldehyde at room temperature for 15 minutes, washed once in PBS containing 0.1% Tween 20 (PBS-T) and permeabilized for 15 minutes in PBS 0.5% Triton X-100. E4.5 embryos were blocked in PBS-T with 2% BSA (Sigma) and 5% normal donkey serum (Jackson ImmunoResearch Laboratories) at room temperature for 2 hours and E6.25 embryos were blocked in PBS-T with 10% BSA (Sigma) and 5% normal donkey serum (Jackson ImmunoResearch Laboratories) at room temperature for 8 hours or overnight at 4°C. Primary and secondary antibodies were diluted in blocking solution. Staining was performed at room temperature for ∼2-5 hours or overnight at 4°C. Washes after primary and secondary antibodies were done three times in PBS-T. In the last washing step (15 minutes) 1:500 Hoechst 33258 (Invitrogen) was used to stain nuclei. E4.5 embryos were mounted in PBS in wells made with Secure Seal spacers (Molecular Probes™, Thermo Fisher Scientific) and placed between two cover glasses for imaging. E6.25 embryos were mounted in agarose plugs using 1% low melting agar (Sigma).

### Immunofluorescent staining of E12.5 placentas

Placentas were dissected from pregnant females at E12.5, washed briefly in ice cold PBS and fixed in 4% PFA overnight. Depending on the experiment, embryos and placentas were prescreened for contribution to individual compartments using a fluorescent stereomicroscope. The following day, placentas were embedded, frozen and sectioned (10μm) starting at the sagittal plane. Sections were blocked with 10% horse serum in PBS for 1-hr at room temperature and incubated overnight at 4°C with primary antibodies diluted with 5% horse serum in PBS. DsRed chimeric placentas were stained with anti-mCherry/Rfp antibody and H2B-Gfp chimeric placentas with anti-GFP. For detection of fetal endothelial cells, placentas were first subjected to antigen retrieval using 10 mM citrate buffer, pH 6.0 at 100°C for 10 minutes, then cooled, blocked, and co-stained with anti-CD34. The following day, sections were stained with the appropriate secondary antibodies for 1-hr at room temperature, washed in PBS, and eventually counterstained with DAPI (Sigma). Sections exposed to secondary antibody alone were used as negative controls. Antibody specificity for mCherry/Rfp and Gfp were confirmed on non-chimeric placentas.

### Antibodies for immunofluorescence stainings

Primary antibodies: rabbit anti-mCherry 1:500 (Ab167453, Abcam) *detects DsRed; mouse anti-mCherry 1:500 (632543, Clontech); chicken anti-Gfp 1:500 (ab13970, Abcam); rabbit anti-Cdx2 1:600 (ab76541, Abcam); rabbit anti-Nanog 1:100 (09-0020, ReproCell); goat anti-Gata3 1:100 (AF2605, RandD Systems), goat anti-Sox2 1:100 (AF2018, RandD Systems); goat anti-Oct4 1:100 (sc-8629, Santa Cruz Biotechnologies); goat anti-Sox17 1:100 (AF1924, RandD Systems); goat anti-Elf5 1:100 (sc9645, SantaCruz); rabbit anti-Tfap2c (sc8977 SantaCruz); rat anti-Krt8 1:10 (TROMA-I antibody, Developmental Studies Hybridoma Bank, Iowa City, IA, USA); and mouse anti Krt18 1:100 (ab668, Abcam). To visualize trophoblast cells in placentas, co-immunostaining was performed with the following antibodies: anti Kr8 (1:10, TROMA-1), anti Tfap2c (1:80, Santa Cruz, sc-8977), anti Cdh3 (1:100, RD, AF761), anti Tpbpa (1:150, Abcam, ab104401), anti Mct1 (1:400, Millipore, AB1286I) and anti Mct4 (1:400, Millipore, AB3314P). Endothelial cells were identified by immunoreactivity to CD34 antibody (1:100, Abcam, ab8158) Secondary antibodies: (diluted 1:500) 448, 549, 594 or 633 conjugated donkey anti-mouse, donkey anti-rabbit or donkey anti-goat DyLight (Jackson ImmunoResearch) or Alexa Fluor (Life Technologies) and donkey anti rat DyLight 488 (Bethyl).

### Imaging

Images of E4.5 embryos were acquired using a Zeiss Axiovert 200 inverted microscope equipped with a Hamamatsu C9100-13 EM-CCD camera, a Quorum spinning disk confocal scan head and Volocity aquisition software (PerkinElmer). Z-stacks were taken at 1μm intervals with a 20x air objective (NA = 0.75). Images of E6.25 embryos were acquired using a Ziess Ligh Sheet Z1, with a pCO Edge 5.5 × 2 camera and Zeiss Zen Lightsheet 2014 acquisition software. Images were acquired using a 20x water immersion detection objective (NA = 1.0). Z-stacks were taken at 1μm intervals. Images of E12.5 placentas were acquired using a Pannoramic 250 Flash II from 3DHistech.

### Data access

All raw and processed sequencing data generated in this study have been submitted to the NCBI Gene Expression Omnibus (GEO; http://www.ncbi.nlm.nih.gov/geo/) under accession number GSE145609.

Processed data can also be visualized and downloaded using .loom files deposited under: https://github.com/pasquelab/totipotency.

## Acknowledgements

The authors wish to acknowledge the contribution of the Model Production Core staff at the Center for Phenogenomics for technical support. Single cell sequencing was performed at National Genomics Infrastructure in Stockholm at Science for Life Laboratory (funded by the Knut and Alice Wallenberg Foundation and the Swedish Research Council) with assistance from SNIC/Uppsala Multidisciplinary Center for Advanced Computational Science with massively parallel sequencing and access to the UPPMAX computational infrastructure. Bulk RNA sequencing was performed in KU Leuven Genomics Core. We would like to thank Stein Aerts, Sara Aibar, Chris Davie and Christopher Flerin for discussions and support. This work was funded by CIHR (FDN-143334), Genome Canada and Ontario Genomics (OGI-099), Programme de bourses de chercheur-boursier FRQS Junior 1 (FRQS 268829, 280187), the Swedish Research Council, Ragnar Söderberg Foundation, Ming Wai Lau Center for Reparative Medicine, Center for Innovative Medicine, Wallenberg Academy Fellow, NSERC (2014-04497). Research in the Pasque lab is supported by The Research Foundation–Flanders (FWO) (Odysseus Return Grant G0F7716N to VP, FWO grants G0C9320N and G0B4420N to VP), the KU Leuven Research Fund (BOFZAP starting grant StG/15/021BF to VP, C1 grant C14/16/077 to VP and project financing), and FWO Ph.D. fellowships to A.Ja (1158318N), IT (1S72719N) and SKT (1S75720N).

## Author contributions

J.P.S., A.M., A.Ja., T.P., M.E.B., N.D.G., S.K.T. performed *in vitro* stem cell experiments. E.P., I.R., B.B. performed *in vivo* chimera experiments. J.P.S., A.Ja., P.K., S.P., M.E.B., I.T., F.L., V.P., performed transcriptional analyses. E.P., V.P., F.L., J.P.S., A.Ju. and J.R., planned experiments, analyzed data and wrote the manuscript. V.P., F.L., E.P., and J.R., conceived the study.

## Supplementary Figures

**Fig. S1: Culture conditions and morphologies of ESCs, L-EPSCs and D-EPSCs** Scale bar 100 µm.

**Fig. S2: Transcriptional profiling of L- and D-EPSCs related to Figure 1**

A) Gene expression comparison of pluripotency and somatic genes between 2iLif ESCs and L-EPSCs and mouse embryonic fibroblasts (MEFs) as control.

B) Changes in expression levels between 2iLif ESCs and L-EPSCs for genes reported to change in Yang et al., 2017^32^.

C) UMAP from Figure 1E showing the resource from which each cell was merged, and lists the number of cells used from each.

D) FeaturePlots projecting expression of representative 2-cell, EPI, PE, and TE marker genes, overlaying Figure1E UMAP.

E) Heatmap of genes previously reported to be upregulated in D-EPSCs compared to ESCs cultured in Lif/serum conditions, which are similarly upregulated in D-EPSCs (generated in this study) when compared to naïve 2iLif ESCs.

F) UMAP constructed based on the activity of gene regulatory networks in 2iLif ESCs, embryonic stages E3.5 ICM, E4.5 EPI, E5.5 EPI, L-EPSCs and D-EPSCs.

**Fig. S3: Re-analysis of transcriptional profiling data of blastoid cells related to Figure 3**

A) UMAP from Figure 3A identifying original datasets for each cell.

B) UMAP from Figure 3A, highlighting ZG-blastoid EPI, ZG-blastoid ZG, B-blastoid TE, and ZG-blastoid intermediate.

C) UMAP from Figure 3A, highlighting ZG-blastoid populations produced with 2iLif vs. EPSCs

D) UMAP from Figure 3A, highlighting B-blastoid EPI, B-blastoid PE, B-blastoid TE, and B-blastoid intermediates.

E) Pie chart showing percent of each cell type category based on our re-analysis of all B-blastoid cells.

**Fig. S4: Gene regulatory network atlas spanning mouse embryo stages from morula to gastrulation and *in vitro* blastoid models**

A) UMAP clustering based on the activity of gene regulatory networks.

B) Heatmap representing the activity of selected regulons associated with lineage-specific TFs averaged across cells from each cluster.

C) Dot plot showing expression levels of NANOG, GATA4, GATA3, T and CDX2 target genes derived from gene regulatory network analysis of selected cell types.

**Fig. S5: E6**.**25 chimeras using mScarlet-NLS ESCs and L-EPSCs**

A-B) Bar charts showing percent of E4.5 and E6.5 chimeric embryos with different lineage contributions. Chimeras were generated by aggregating a wild-type host E2.5 embryo with a small clump of mScarlet ESCs (129 ESC, R1-ES mScarlet-NLS) which were cultured in either 2iLif or L-EPSC conditions. n indicates number of chimeras generated.

C) Immunofluorescent staining of E4.5 chimeric embryo stained for mCherry (mScarlet), Sox2 (EPI) and Cdx2 (TE). Donor cells were cultured in 2iLif conditions. Scale bar = 20 µm.

C’) Magnified single plane showing mCherry/Sox2 positive, Cdx2 negative cell within the TE. Scale bar = 20 µm.

D) Immunofluorescent staining of E4.5 chimeric embryo. Donor cells were cultured in L-EPSC conditions. Merges show a magnified single plane of the ICM. Scale bar = 20 µm.

E) Immunofluorescent staining of E6.5 chimeric embryo cultured in 2iLif conditions stained for mCherry (mScarlet) Oct4 (EPI). Maximum intensity projection. Scale bar = 20 µm.

E’) Magnified single plane showing single mCherry/Oct4 positive donor cell found within the trophoblast compartment. Scale bar = 20 µm.

F) Immunofluorescent staining of E6.5 chimeric embryo stained for mCherry (mScarlet) and Elf5 (trophoblast compartment). Donor cells were cultured in L-EPSC conditions. Single plane image. Scale bar = 20 µm.

F’) Magnified single plane showing single donor mCherry positive, Elf5 negative cell found within the epiblast region. Scale bar = 20 µm.

**Fig. S6: Analysis of chimeric E12**.**5 placentas generated with totipotent blastomeres** Immunofluorescent stainings of E12.5 chimeric placentas generated with a single blastomere of an 8-cell stage embryo that expressed either H2B-Gfp or DsRed and a wild-type host embryo. H2B-Gfp expressing placentas were stained for Tfap2c, Tpbpa, Mct4, Cd31 and Gfp (top panel). DsRed expressing placentas were stained for Krt8, Mct1, Cdh3 and mCherry/Rfp (bottom panel). Images shown at different magnifications. Scale bars: 500 µm (4x), 200 µm (7x), 50 µm (40x).

**Fig. S7: Full panel of chimeric E12**.**5 placentas generated using tetraploid complementation**

Immunofluorescent staining of E12.5 chimeric placentas generated with H2B-Gfp expressing tetraploid host embryo and wild-type ESCs (left panel), wild-type tetraploid host embryo and H2B-Gfp expressing ESCs (middle panel) and wild-type host embryo (diploid) and H2B-Gfp expressing L-EPSCs. Placenta sections are immuno-stained for Gfp and the following trophoblast markers: Cdh3, Tpbpa, and Mct4, as well as the fetal endothelial cell marker Cd31. A total of 14 H2B-Gfp positive L-EPSCs placentas were collected from two females, and 3 to 4 placentas were analyzed for each marker. Images shown at different magnifications. Scale bars: 500 µm (4x), 200 µm (7x), 50 µm (40x).

