## Supplementary figures and images for "Defining totipotency using criteria of increasing stringency"

### Figure S1

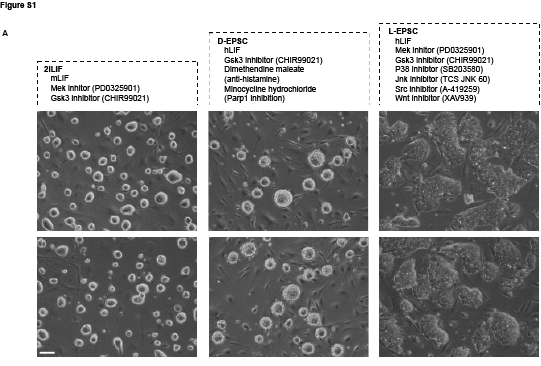

### Figure S2

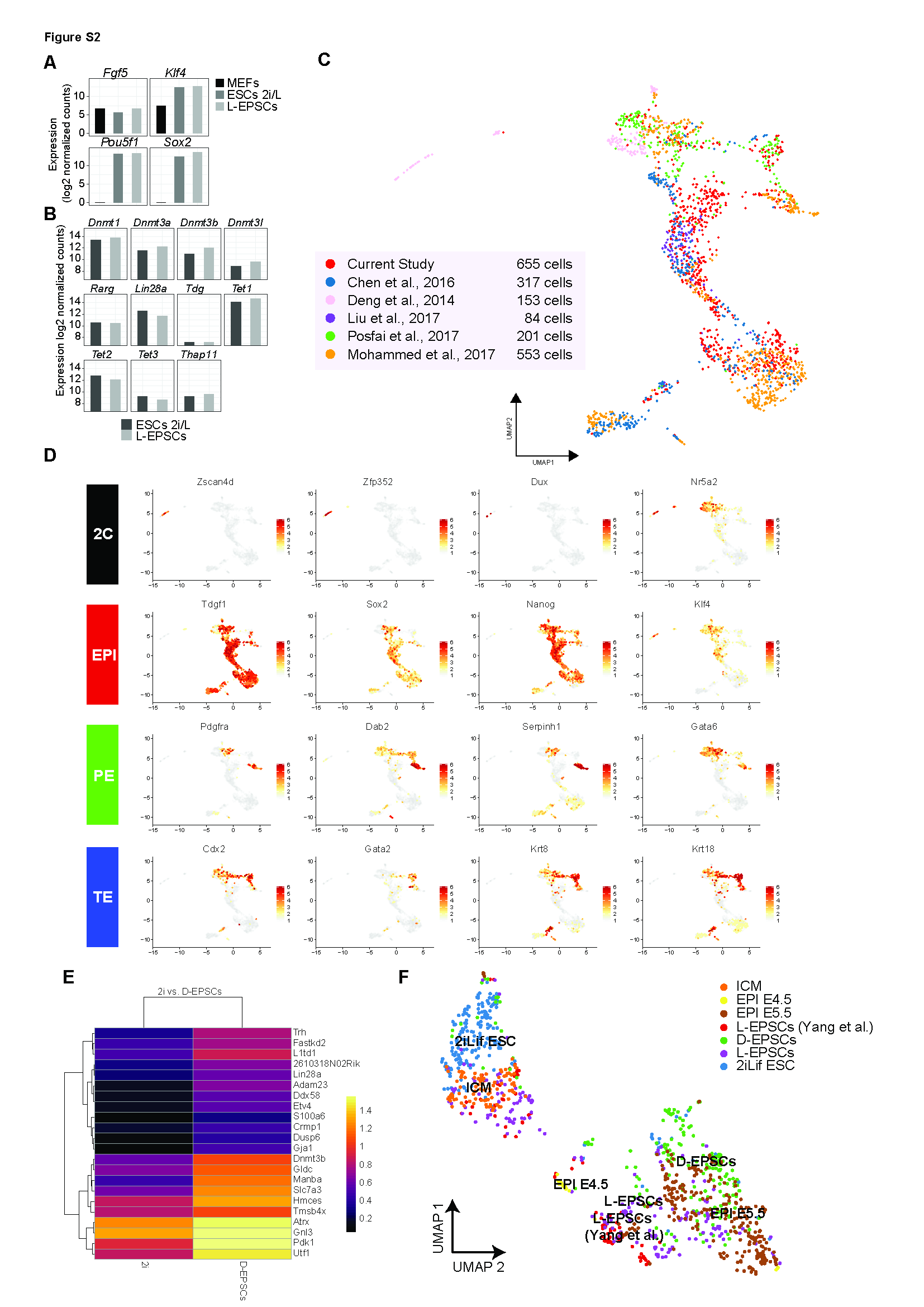

### Figure S3

Figure S3

A

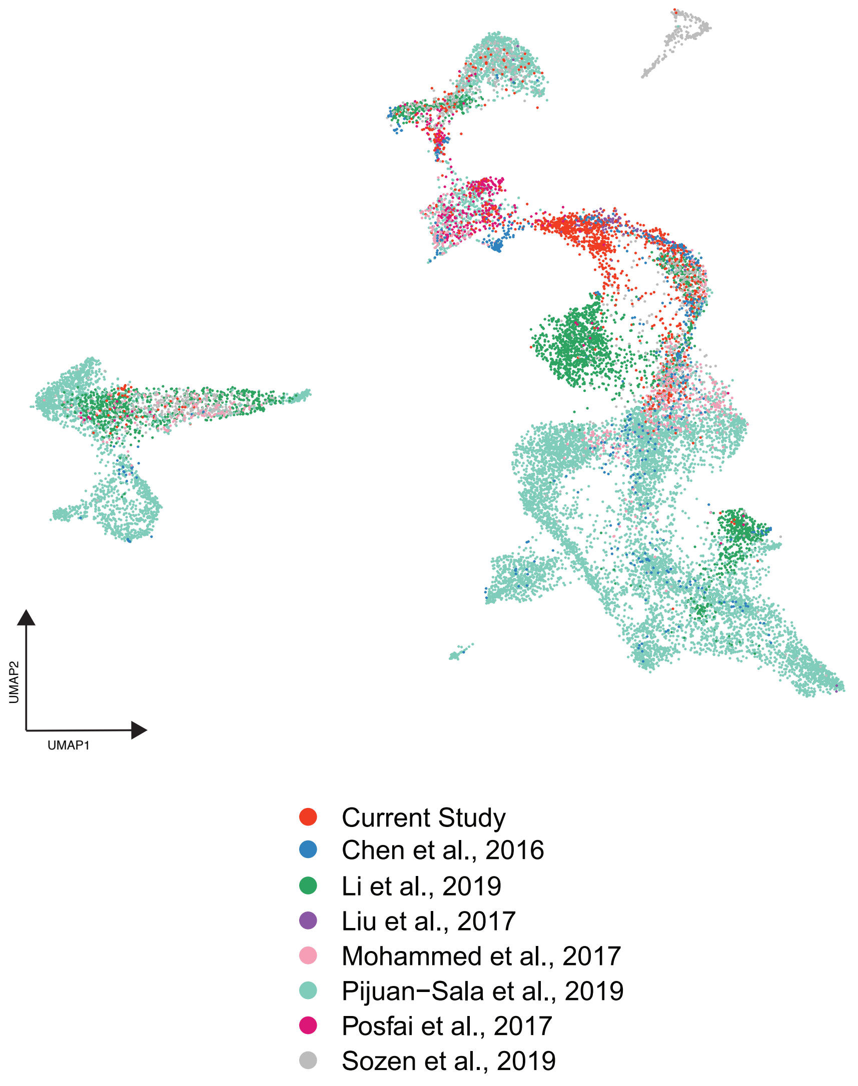


B

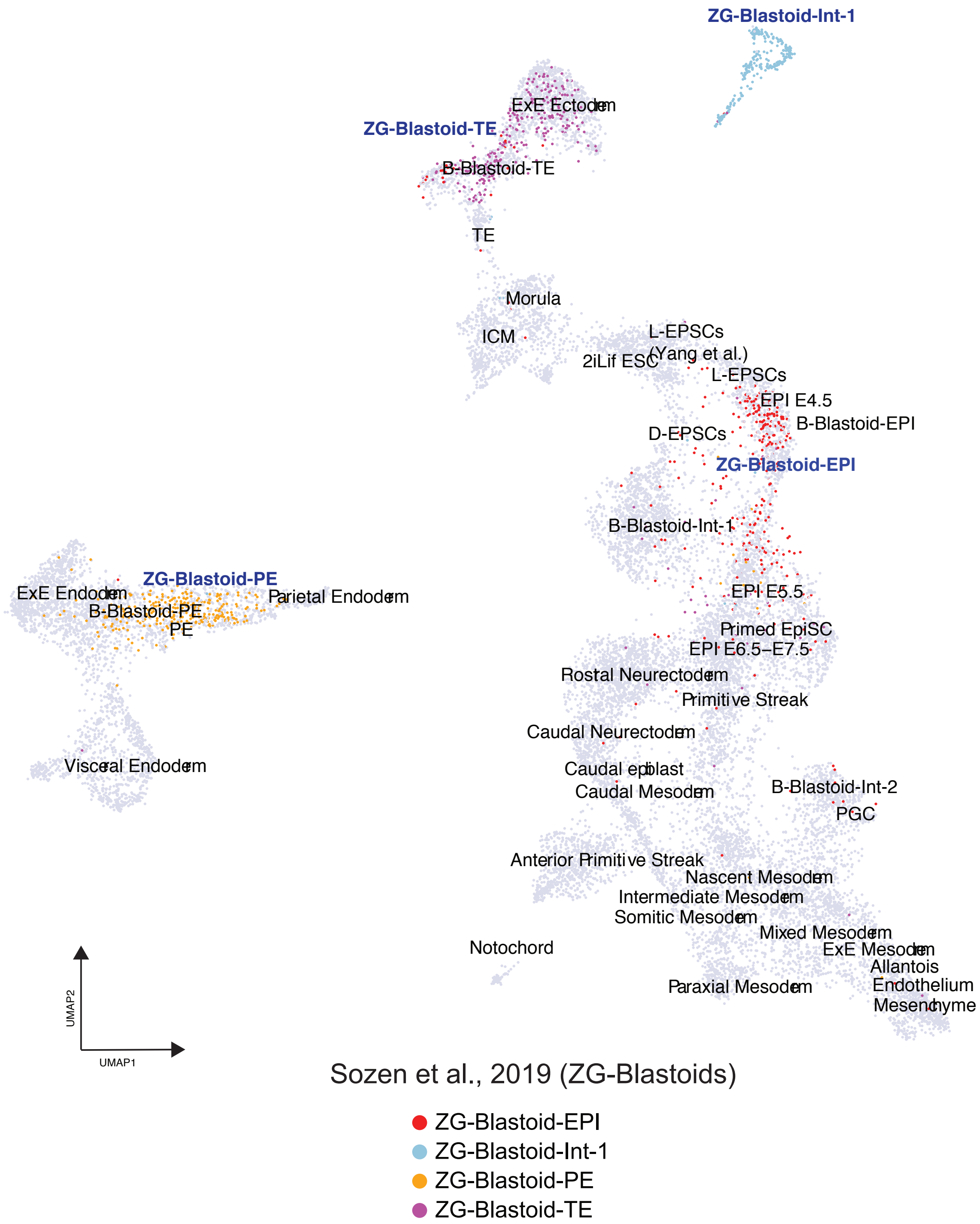


C

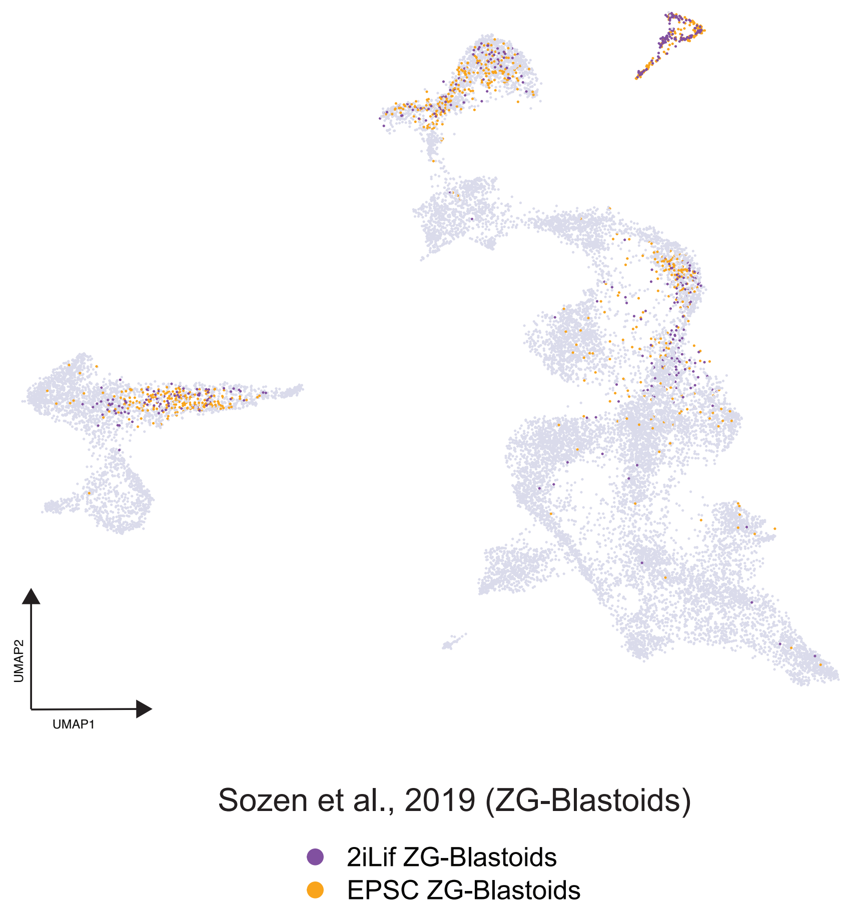


D

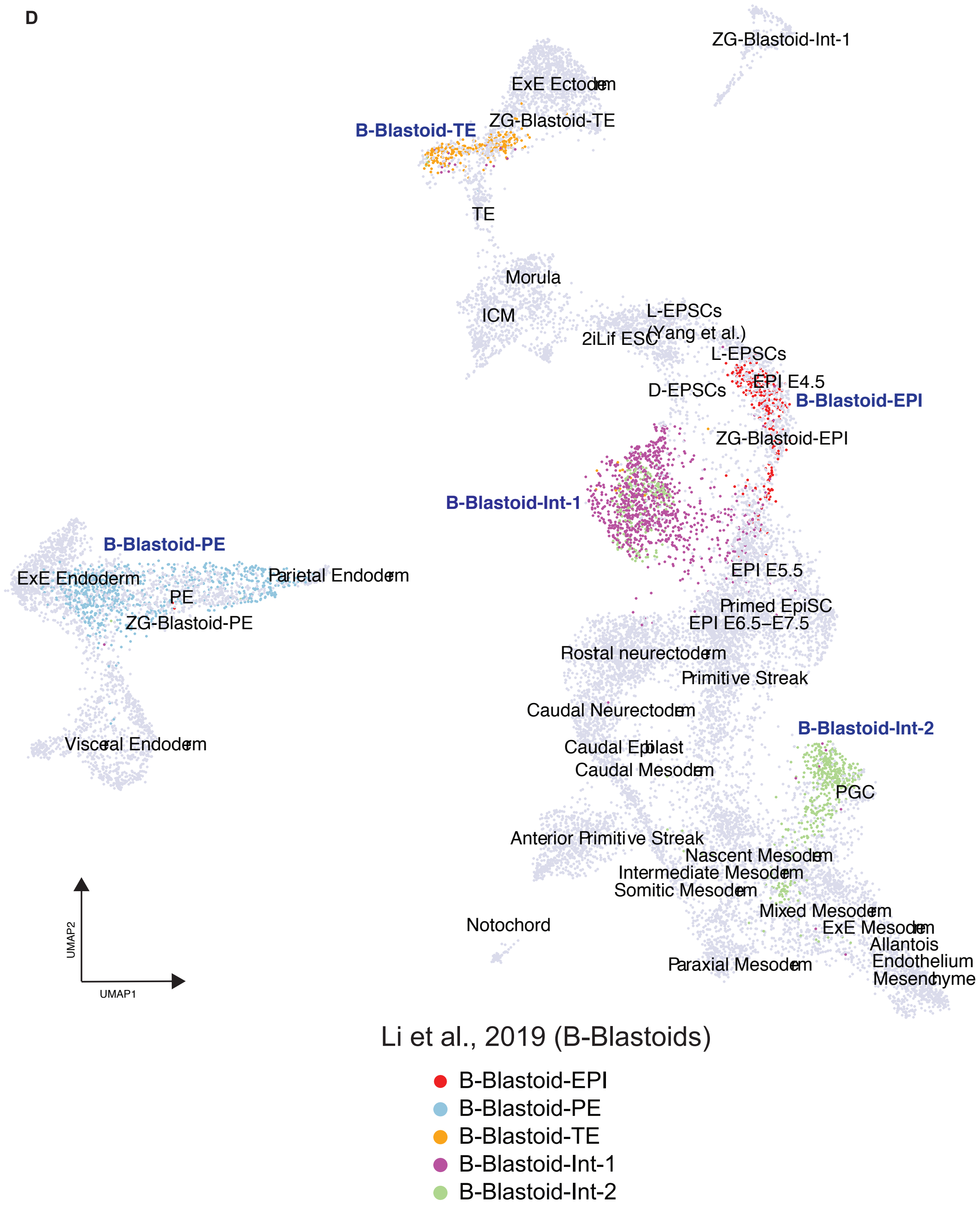


E

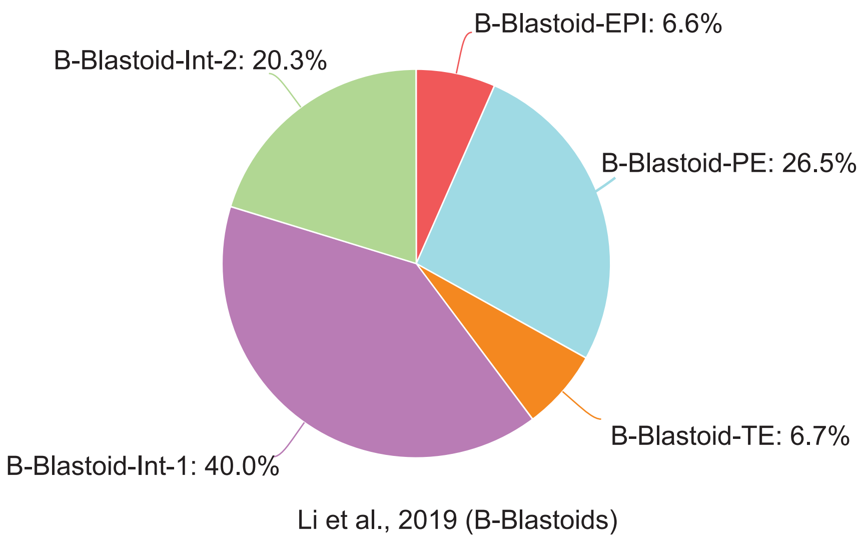

### Figure S4

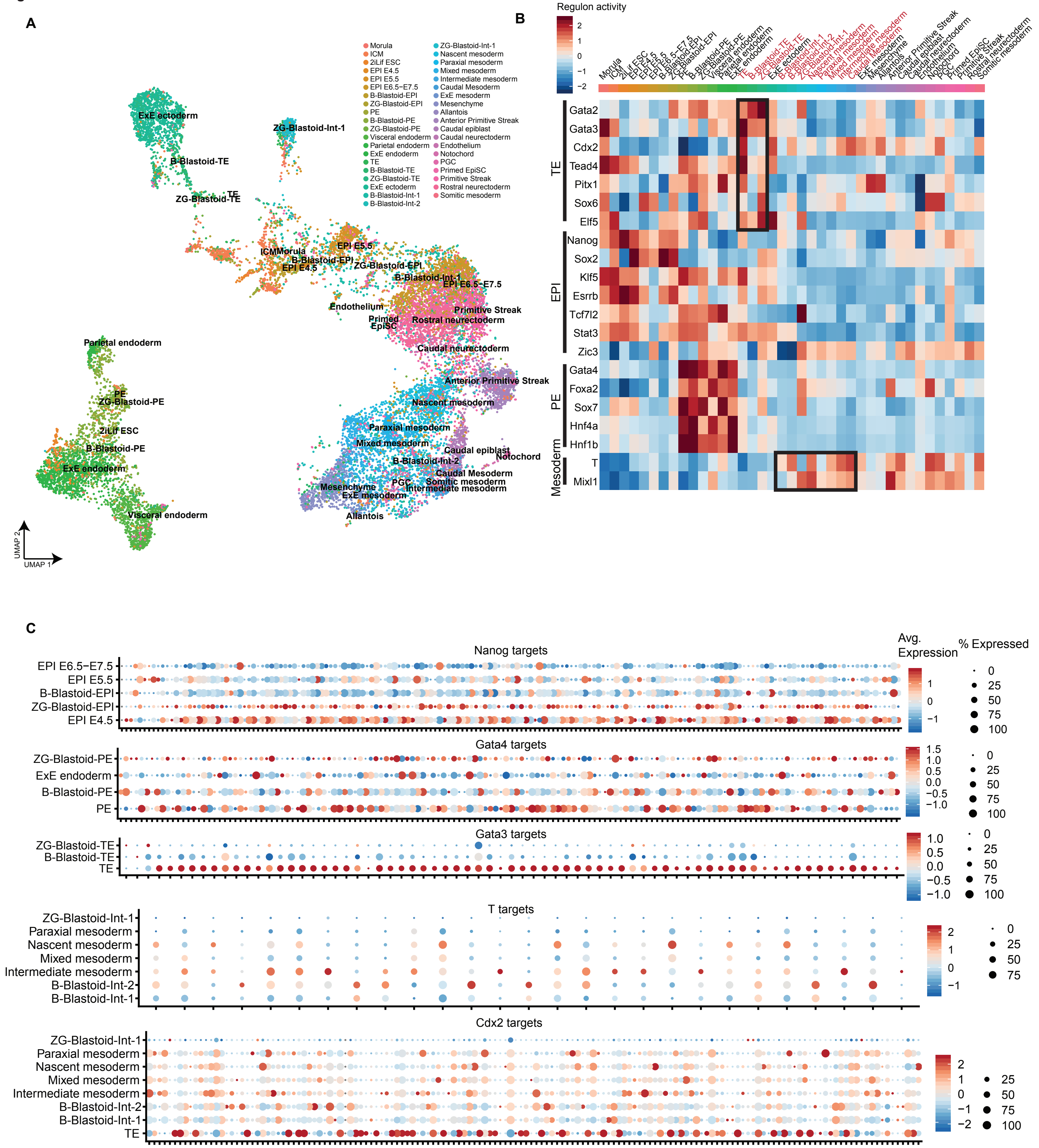

### Figure S5

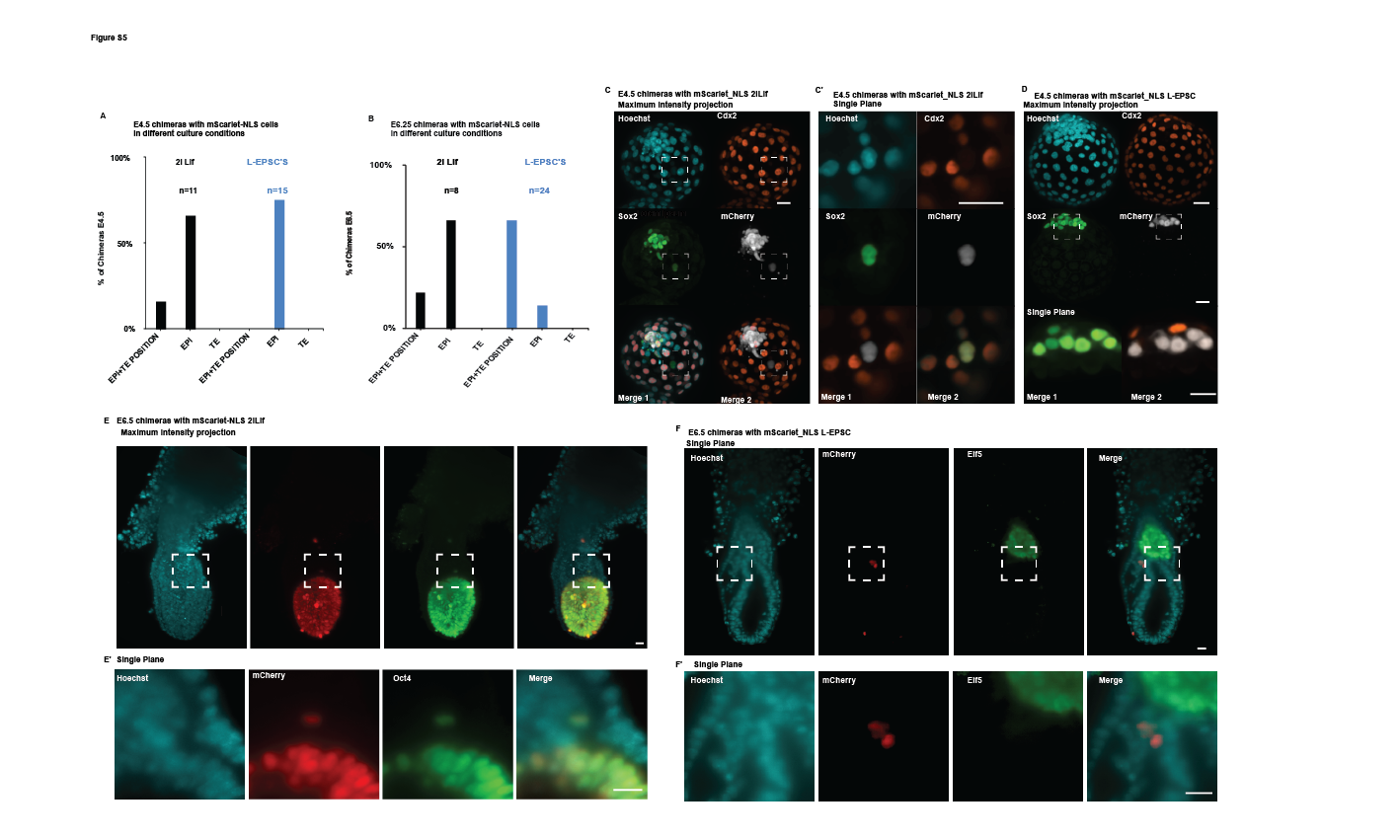

### Figure S7

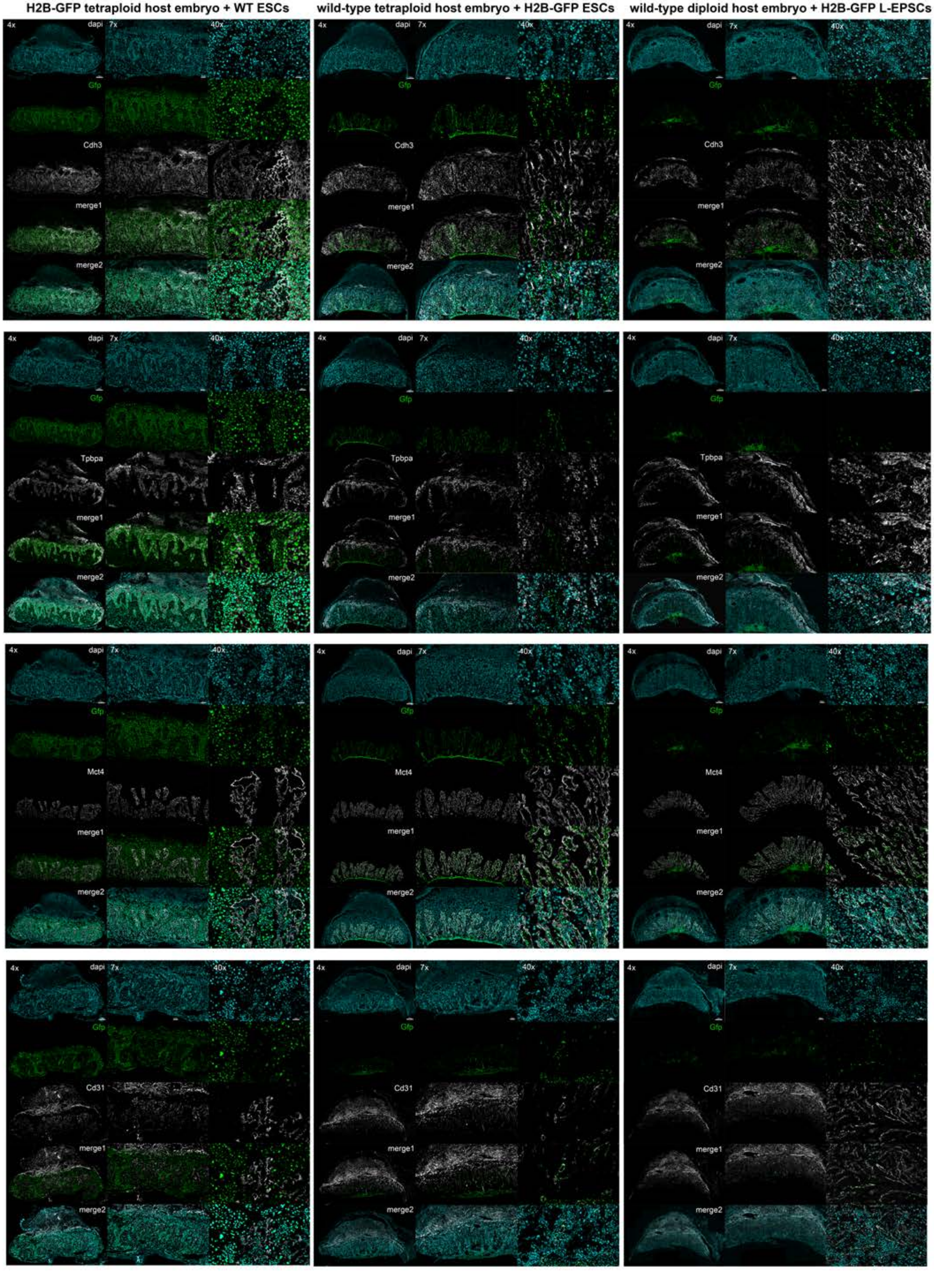
