## Supplementary material for "Defining totipotency using criteria of increasing stringency": Figure S6

**wild-type diploid host embryo + totipotent H2B-Gfp blastomere**

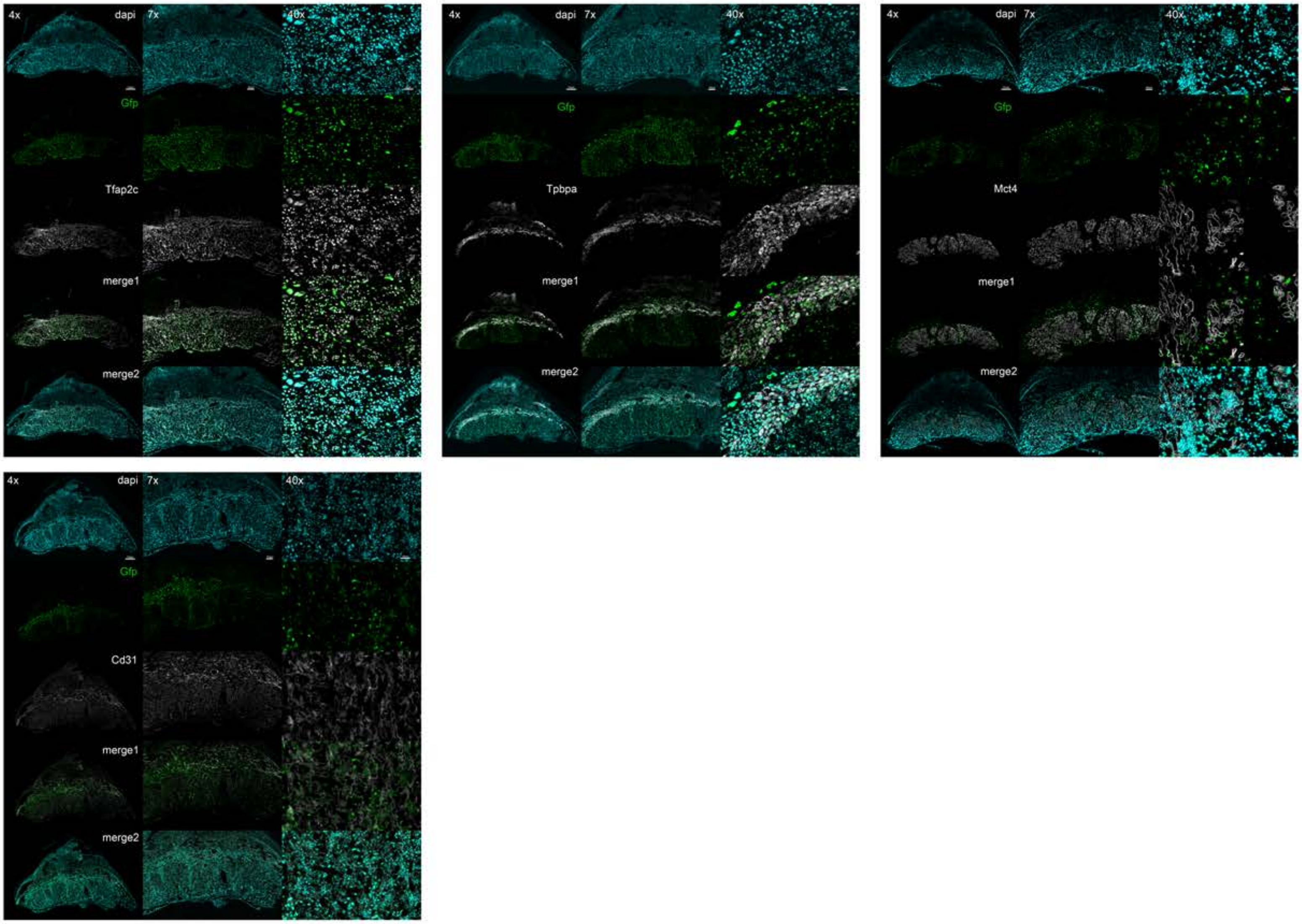


**wild-type diploid host embryo + totipotent DsRed blastomere**

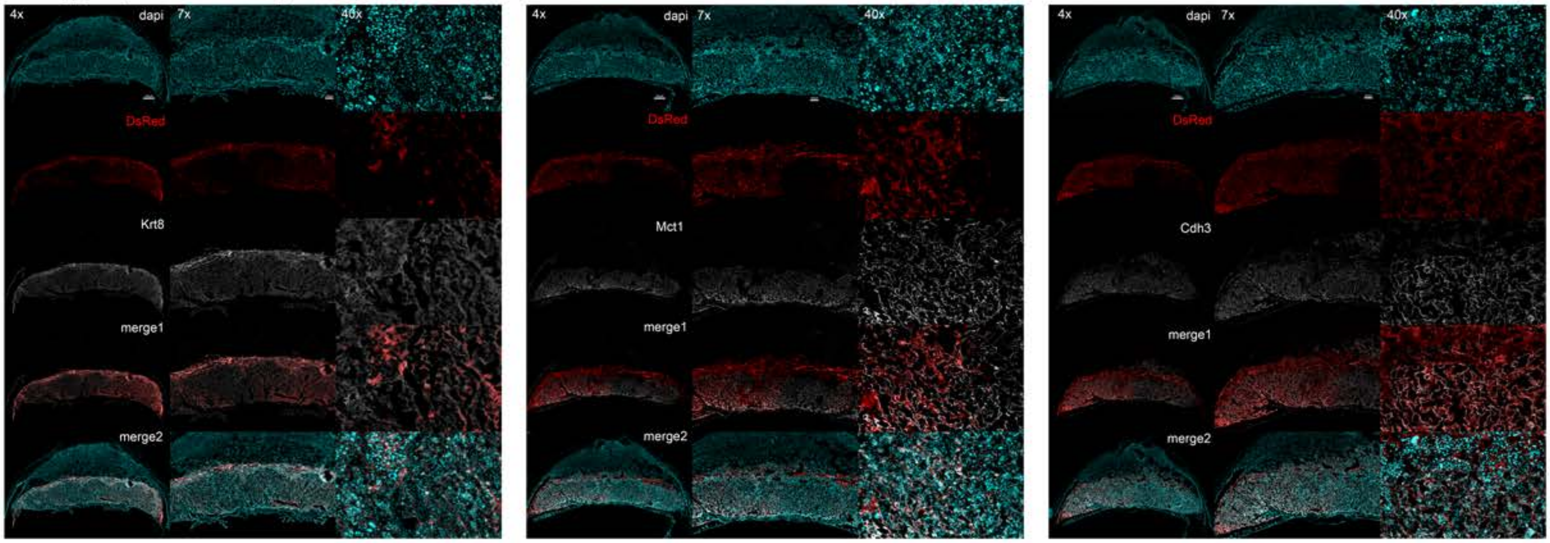
